# Dissection of central clock function in *Drosophila* through cell-specific CRISPR-mediated clock gene disruption

**DOI:** 10.1101/640011

**Authors:** Rebecca Delventhal, Meghan Pantalia, Reed M. O’Connor, Matthew Ulgherait, Han X. Kim, Maylis Basturk, Julie C. Canman, Mimi Shirasu-Hiza

## Abstract

In *Drosophila*, ~150 neurons expressing molecular clock proteins regulate circadian behavior. Sixteen of these clock neurons secrete the neuropeptide Pdf and have been called “master pacemakers” because they are essential for circadian rhythms. A subset of Pdf^+^ neurons (the morning oscillator) regulates morning activity and communicates with other non-Pdf^+^ neurons, including a subset called the evening oscillator. It is assumed that the molecular clock in Pdf^+^ neurons is required for these functions. To test this, we developed and validated Gal4-UAS based CRISPR tools for cell-specific disruption of key molecular clock components, *period* and *timeless*. While loss of the molecular clock in both the morning and evening oscillators eliminates circadian locomotor activity, the molecular clock in either oscillator alone is sufficient for circadian locomotor activity. This suggests that clock neurons do not act in a hierarchy but as a distributed network to regulate circadian activity.

## Introduction

Circadian rhythms are 24-hour oscillations in physiological functions and behaviors, including locomotor activity, immune system function, metabolism, and sleep (1–7). Disruption in circadian regulation is a common feature of aging and is associated with a variety of adverse health outcomes such as diabetes and cancer (8–11). Circadian rhythms are driven by “molecular clocks,” or proteins that regulate rhythmic gene expression. Work in *Drosophila* has been crucial for understanding the molecular clock, a transcriptional negative feedback loop with four core proteins: Clock, Cycle, Period, and Timeless (Fig. 1A) (12–17). In brief, Clock and Cycle activate transcription of *period* and *timeless* which, once translated, dimerize and translocate into the nucleus where they bind to Clock and Cycle, thereby inhibiting their own transcription; this molecular feedback loop repeats with a 24-hour periodicity (Fig. 1A). Importantly, the core components of the molecular clock in *Drosophila* are conserved in humans (18).

**Figure 1.**
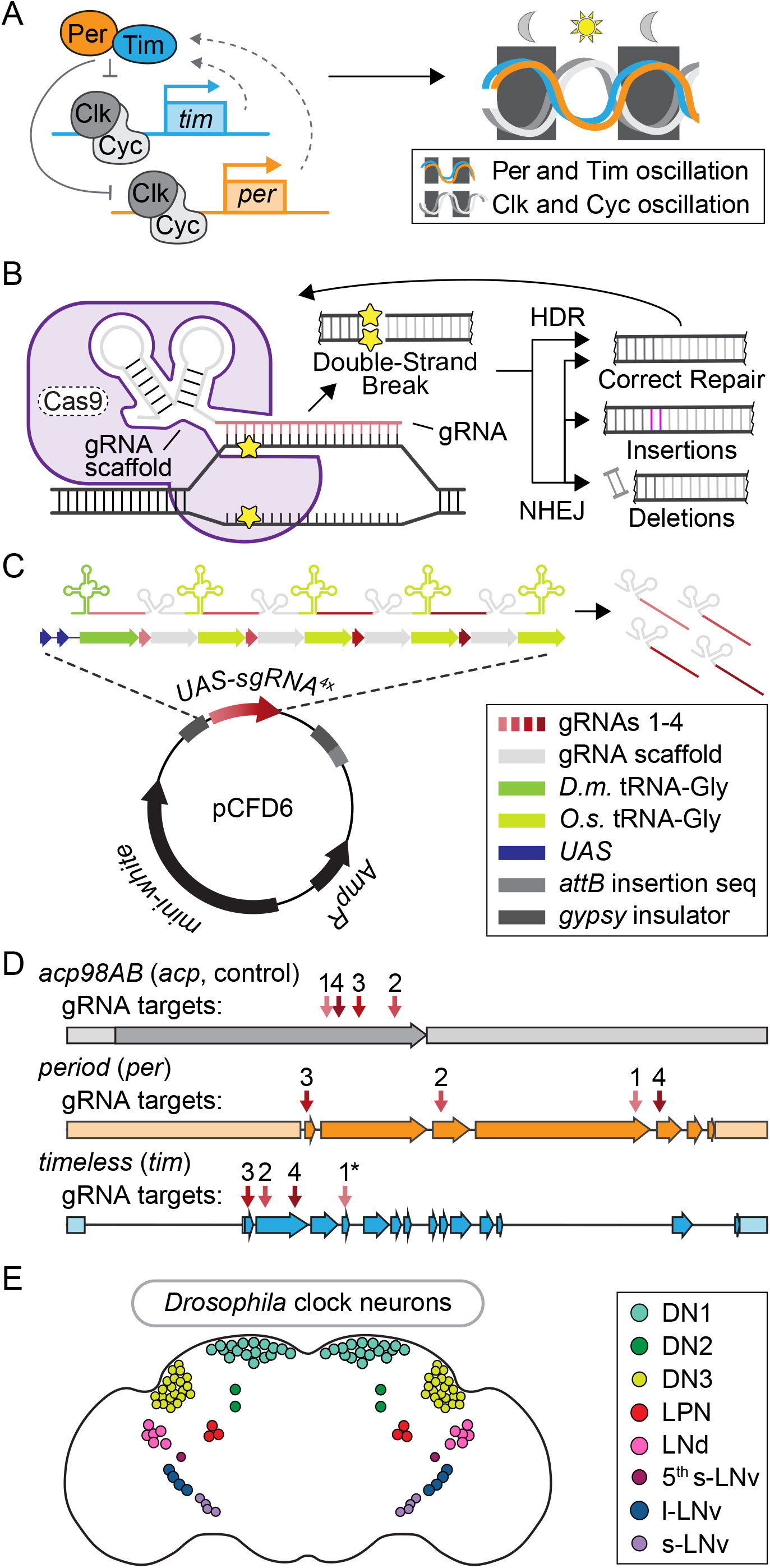
Toolbox for cell-specific, CRISPR-mediated disruption of core circadian regulators. (A) Schematic of the transcriptional/translational negative feedback loop that drives rhythmic expression and activity of the four core circadian regulators: Period (Per), Timeless (Tim), Clock (Clk), and Cycle (Cyc). (B) Diagram of CRISPR-Cas9 mediated DNA damage and repair pathways. (C) Diagram of plasmid (pCFD6, adapted from Port and Bullock 2016 (35)) used to generate *UAS-sgRNA*^*4x*^ transgenic flies. *D.m*. = *Drosophila melanogaster*. *O.s.* = *Oryza sativa*, Asian rice. (D) Diagram showing sgRNA target sites for *acp98AB* (*acp*, gray), *period* (*per*, orange), and *timeless* (*tim*, blue), numbered in order of 5’-3’ position in the respective *UAS-sgRNA*^*4x*^ construct. Arrows = exons; shaded rectangles = promoters and UTRs. \**tim* sgRNA 1 has a single base pair deletion in the Cas9-binding scaffold region (see Methods). (E) Diagram of ~150 clock neurons organized into the following anatomical and functional clusters in the *Drosophila* brain: dorsal neurons (DN1, DN2, DN3), lateral posterior neurons (LPN), dorsal lateral neurons (LNd), and small and large ventral lateral neurons (s-LNv, 5^th^ s-LNv, l-LNv).

In *Drosophila*, ~150 neurons in the brain have molecular clocks and control circadian locomotor activity (Fig. 1E) (19). These clock neurons cluster in eight subgroups defined by their anatomical locations: small and large ventral lateral neurons (s-LNvs and l-LNvs), the 5^th^ s-LNv, dorsal lateral neurons (LNds), lateral posterior neurons (LPNs), and three separate clusters of dorsal neurons (DN1s, DN2s, and DN3s) (Fig. 1E). Cell ablation and cell-specific rescue experiments identified two sets of clock neurons that control circadian locomotor activity: Pdf^+^ s-LNvs comprise the “morning oscillator” and control the morning peak of activity, while the 5^th^ s-LNv and LNds comprise the “evening oscillator” and control the evening peak of activity (20–22). In the classic paradigm of circadian neuronal circuitry, the morning oscillator neurons are thought to be master regulatory neurons that synchronize molecular clocks in other neurons via rhythmic release of the neuropeptide Pigment-dispersing factor (Pdf) (19, 23–26). However, a subset of *Pdf* mutants (~25%) were reported to retain rhythmic activity with a shortened period (21) and more recent experiments involving cell-specific expression of period-lengthening and shortening genes have suggested that circadian neurons interact through a complex network, rather than a hierarchy, to regulate circadian behavior (22, 27). The precise role of molecular clock components in these circadian-regulatory neurons remains unclear.

To assess the role of molecular clock components in specific clock neurons, researchers have typically used the Gal4-UAS system for cell-specific RNAi-knockdown of clock genes and cell-specific rescue in a null mutant (28, 29). While instrumental in understanding neuronal control of circadian behaviors, these strategies have limitations. RNAi can be inefficient: Martinek and Young observed only ~50% reduction in *per* RNA levels with eye-specific RNAi knockdown of *per* (28). Moreover, unlike *per* null mutants, which are 100% arrhythmic, flies with *per* RNAi knockdown in all Tim^+^ cells were shown to be only 45% arrhythmic (30) or rhythmic with lengthened period (28). Similarly, cell-specific rescue experiments sometimes do not reproduce wild-type rhythmic behavior, possibly due to constitutive expression of normally rhythmic genes. Pan-neuronal or ubiquitous rescue of *per* or *tim* in a null mutant background caused variable rhythmicity (~50-95%), depending on the UAS transgene insertion and Gal4 driver lines used; even overexpression of *per* and *tim* in a wild-type background sometimes resulted in a partial loss of rhythmicity (31). Thus, while cell ablation experiments have shown the necessity of specific neurons for regulation of circadian locomotor activity, the function of the molecular clock within those neurons remains unclear.

Recent advances in CRISPR technology in *Drosophila* provided an opportunity to create new tools for circadian research (32, 33). One key advance was the generation of loss of function (LOF) mutations in somatic cells via biallelic gene-targeting, using UAS-driven expression of the Cas9 enzyme under Gal4 control (34). Briefly, an sgRNA (Cas9 scaffold plus guide RNA) directs Cas9 to the complementary target DNA sequence and catalyzes a double-strand break (DSB) (Fig. 1B). Repair of this DSB occurs either by precise homology-directed repair (HDR) or more error-prone non-homologous end joining (NHEJ) (Fig. 1B). If the targeted sequence is repaired correctly, it will be targeted by the CRISPR machinery for DSB again. If it is repaired incorrectly, this could result in small insertions or deletions (Fig. 1B), which can cause frame-shift mutations, early stop codons, and loss of function (34). Additionally, placing tRNA sequences between multiple sgRNAs in a single transcript allows their release by endogenous tRNA excision machinery and improves the efficiency of gene disruption (35). For example, when Port and Bullock used this strategy to express four unique sgRNAs together, ~100% of the eye area exhibited the LOF *sepia* phenotype, compared with only 11-58% from each individual sgRNA expressed alone. Thus, targeting multiple unique sgRNAs to the same gene increases the likelihood of achieving a LOF mutation in that gene (35). Finally, expressing both the Cas9 enzyme and the sgRNA sequences from two separate UAS-transgenes reduced gene disruption in non-target tissues, likely due to the low probability of having sufficiently leaky expression of both UAS transgenes without a Gal4 present (35).

Here we generated *UAS* transgenes expressing multiple sgRNAs that target either *timeless*, *period*, or a control gene (*acp*). We validated these constructs by showing that CRISPR-mediated gene disruption of *tim* or *per* recapitulates null mutant phenotypes when driven in all clock neurons (Tim^+^ cells), but not in glia, and further confirmed gene disruption by qRT-PCR over the circadian cycle and brain immunostaining. We then disrupted the molecular clock in both the morning and evening oscillators (Mai179^+^), only in the morning oscillator (Pdf^+^), or only in the evening oscillator (Mai179^+^Pdf^−^). These experiments showed that, in Pdf^+^ neurons (which include the morning oscillator), the molecular clock is not necessary, but is sufficient for circadian locomotor activity, challenging the assumption that these neurons require an internal molecular clock to synchronize the activity of other clock neurons. This further suggests that circadian neurons act in a distributed network that can compensate for loss of the molecular clock in specific subsets.

## Results

### *UAS-sgRNA* constructs target circadian gene expression in a tissue-specific manner

We generated CRISPR tools for cell-specific gene disruption of *period* (*per*) and *timeless (tim)* (Fig. 1A), based on previous work (34, 35). UAS-driven constructs with multiple scaffold-guide RNAs (sgRNAs) were paired with a *Gal4* expression driver and a *UAS-Cas9* construct to induce cell-specific LOF mutations (Fig. 1B). We refer to this combination of *Gal4*-driven *UAS-sgRNA* and *UAS-Cas9* expression as “(*target gene)*^*CRISPR*^”. In addition to *tim* and *per*, we also targeted the control gene *acp98AB* (*acp*). Because *acp* is expressed exclusively in male accessory gland cells and the testes (36, 37), CRISPR-mediated mutation of this gene in neurons serves as a control for any nonspecific effects due to double-strand DNA break events, such as cell death. To clone the *UAS-sgRNA* lines, we used the *pCFD6* plasmid designed by Port & Bullock (Fig. 1C) (35). The cassette contains four unique gRNA sequences (see Methods) that target the first four exons of the gene of interest to ensure efficient and specific gene disruption (Fig. 1D).

To determine which circadian neurons require *per* and *tim* expression to influence behavioral rhythmicity, we used three previously characterized *Gal4* drivers that express in clock neurons. *Tim-Gal4* drives expression in all clock gene-expressing cells in the body, including all ~150 clock neurons (38) (Fig. 1E). *Mai179-Gal4* drives expression in a distinct subset of clock neurons that include both morning and evening oscillator neurons: s-LNvs, 5^th^ s-LNv and 3 CRY^+^ LNds with weak and variable expression in DN1s and l-LNvs (39). *Pdf-Gal4* drives expression in the s- and l-LNvs, which express the circadian neurotransmitter *Pdf* (40) and include the morning oscillator (20, 21, 41).

### CRISPR-mediated disruption of *per* or *tim* in all *tim*-expressing cells causes complete loss of behavioral and molecular rhythmicity

To test our *UAS-sgRNA* constructs, we expressed each with *UAS-Cas9* in all Tim^+^ cells using the *tim-Gal4* driver and measured circadian locomotor activity. Flies were entrained in light/dark (LD) conditions and then shifted to constant darkness (DD) to monitor endogenous circadian locomotor activity. We found that CRISPR-targeting *per* or *tim* in Tim^+^ cells (*tim-Gal4*>*per*^*CRISPR*^ or *tim-Gal4*>*tim*^*CRISPR*^) led to complete loss of rhythmic behavior in DD (0% rhythmicity) (Fig. 2A-D). Thus, CRISPR-mediated disruption of *tim* or *per* in Tim^+^ cells, which includes all clock neurons, faithfully recapitulated *tim* and *per* null mutant phenotypes (15, 42). Control flies (*tim-Gal4*>*acp*^*CRISPR*^) maintained circadian locomotor activity (94% rhythmic, 24.46 hr period; Fig. 2B, C), indicating that nonspecific effects from UAS-Cas9 expression or CRISPR-induced double stranded breaks in Tim^+^ cells do not cause loss of rhythms. Together, our results suggest that CRISPR-targeting of *tim* and *per* results in complete functional ablation of the molecular clock, in contrast to the lengthened rhythms sometimes observed with RNAi, which are thought to reflect incomplete reduction of gene expression (28, 43).

**Figure 2.**
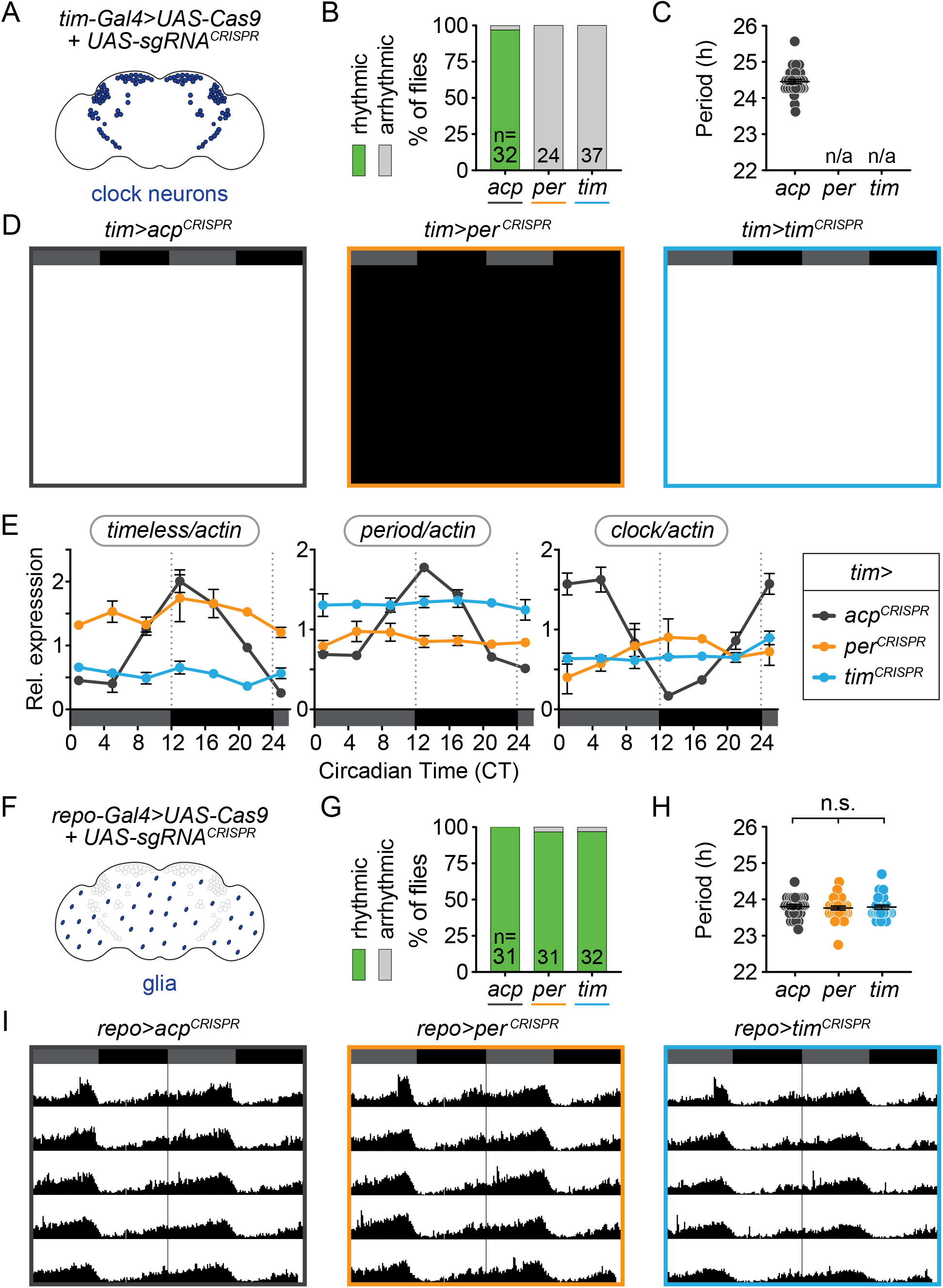
Cell-specific disruption of *per* or *tim* in circadian (Tim^+^) cells but not glial (Repo^+^) cells causes loss of behavioral rhythmicity. (A) Diagram of clock neurons targeted for CRISPR-mediated gene disruption using *tim-Gal4*. (B) Disruption of *per* or *tim* in all clock neurons caused complete loss of behavioral rhythmicity. (C) Scatter plot showing the period of rhythmic flies with *tim-Gal4*-driven disruption of *acp*, *per*, or *tim*. (D) Actograms showing average activity in constant darkness of flies with *tim-Gal4*-driven disruption of *acp*, *per*, or *tim*. Activity data is double-plotted, six days of activity are displayed. Dark grey rectangles = subjective day, black rectangles = subjective night. (E) Relative mRNA levels, measured by qRT-PCR over a 24-hour period, of the core circadian genes *timeless* (left), *period* (middle), and *clock* (right) in heads of *tim-Gal4* CRISPR flies. (F) *repo-Gal4* targets all glia for CRISPR-mediated gene disruption. (G) *repo-Gal4*-driven, CRISPR-mediated gene disruption in glia had no effect on behavioral rhythmicity. (H) Scatter plot showing the period of rhythmic flies with *repo-Gal4*-driven disruption of *acp*, *per*, or *tim*. (I) Actograms show average activity of flies in constant darkness with *repo-Gal4*-driven disruption of *acp*, *per*, or *tim* in glia.

To confirm that rhythmic transcription of circadian clock genes is disrupted by CRISPR-targeting *tim* or *per* in Tim^+^ cells, we analyzed mRNA from fly heads collected over the circadian cycle. In wild-type fly heads, clock gene mRNA levels oscillate with approximately 24-hour periodicity in constant darkness (15, 16, 44). We found that control flies (*tim-Gal4*>*acp*^*CRISPR*^) also displayed robust oscillation of *timeless*, *period*, and *clock* transcripts (Fig. 2E, grey). In contrast, CRISPR-targeting of *tim* or *per* in Tim^+^ cells resulted in arrhythmic transcription of all three molecular clock genes (Fig. 2E, green and blue), consistent with the behavioral arrhythmicity caused by these manipulations (Fig. 2B). Furthermore, *tim* transcript levels in *tim-Gal4*>*tim*^*CRISPR*^ flies and *per* transcript levels in *tim-Gal4*>*per*^*CRISPR*^ flies were reduced to levels similar to the lowest baseline levels for these transcripts in control flies. We note that *tim* transcripts, though arrhythmic, were elevated after disruption of *per* (*tim-Gal4*>*per*^*CRISPR*^ flies) and vice versa for *per* transcripts after disruption of *tim*. These results are consistent with earlier findings indicating that loss of either Per or Tim, inhibitors of Clock/Cycle, yields constitutive activity of the Clock/Cycle transcription complex and elevated levels of *per* or *tim* transcripts (12, 45).

To further confirm the efficiency of our gene disruption, we performed immunofluorescence analysis on the brains of CRISPR-targeted flies (*tim-Gal4*>*gene*^*CRISPR*^) for Per and Tim at ZT0, along with *per*^01^ null mutants (Fig. S1A). At ZT0, Per and Tim proteins are highly expressed and localized to the nucleus in wild-type flies (17, 45, 46). Consistent with this, control flies (*tim-Gal4*>*acp*^*CRISPR*^) exhibited high levels of Per and Tim protein expression, colocalized in the nucleus. In contrast, in flies CRISPR-targeted for *per* or *tim* in Tim^+^ cells (*tim-Gal4*>*per*^*CRISPR*^ and *tim-Gal4*>*tim*^*CRISPR*^), we observed loss of nuclear Per or Tim staining, similar to *per*^01^ null mutants (Fig. S1A). CRISPR-targeting of *per* led to loss of Per signal and only cytoplasmic Tim signal; CRISPR-targeting of *tim* led to loss of both Tim and Per signal, presumably because Per is unstable without Tim (Fig. S1B) (46, 47). Taken together, these results show that CRISPR-mediated*, Gal4-*driven disruption of *per* and *tim* in Tim^+^ cells is highly efficient on both the mRNA and protein levels and is sufficient to block locomotor activity rhythms.

### CRISPR-mediated disruption of *per* or *tim* in glia (Repo^+^ cells) does not disrupt behavioral rhythmicity

As a second control for the effect of CRISPR-induced DNA damage and to confirm that this CRISPR gene targeting is Gal4-specific, we CRISPR-targeted *tim* and *per* in glia, using the pan-glial driver *repo-Gal4* (48, 49). Glia are not predicted to control circadian locomotor activity via circadian clock gene expression (30). We found that nearly all flies, whether CRISPR-targeted for *tim*, *per*, or *acp*, were highly rhythmic (100% of *repo-Gal4*>*tim*^*CRISPR*^ and 97% of *repo-Gal4*>*per*^*CRISPR*^ and *acp*^*CRISPR*^) (Fig. 2F-I). These results demonstrate that there is no leaky or non-Gal4-mediated expression of both the *UAS-Cas9* and *UAS-sgRNA* that affects rhythmicity. These results further confirm previously published results that, while circadian locomotor activity requires intact glial cells, it does not require glial expression of clock genes (30).

### Disruption of *per* or *tim* in both morning and evening oscillators (Mai179^+^ neurons) causes complete loss of circadian locomotor activity

To test the effect of disrupting *per* or *tim* in both the morning and evening oscillators, we expressed our *UAS-sgRNA* constructs in the s-LNvs (including the 5^th^ s-LNv) and 3 CRY^+^ LNds, with weak or variable expression in l-LNvs, DN1s, and non-clock neurons, using the *Mai179-Gal4* driver (Fig. 3A) (21, 22, 39, 50). We found that 100% of flies CRISPR-targeted for *per* and *tim* in *Mai179*^+^ cells (*Mai179-Gal4*>*per*^*CRISPR*^ and *Mai179-Gal4*>*tim*^*CRISPR*^) were arrhythmic, while 91% of control flies (*Mai179-Gal4*>*acp*^*CRISPR*^) remained rhythmic (Fig. 3A-D). Thus, the molecular clock is completely required in Mai179^+^ neurons for circadian locomotor activity.

**Figure 3.**
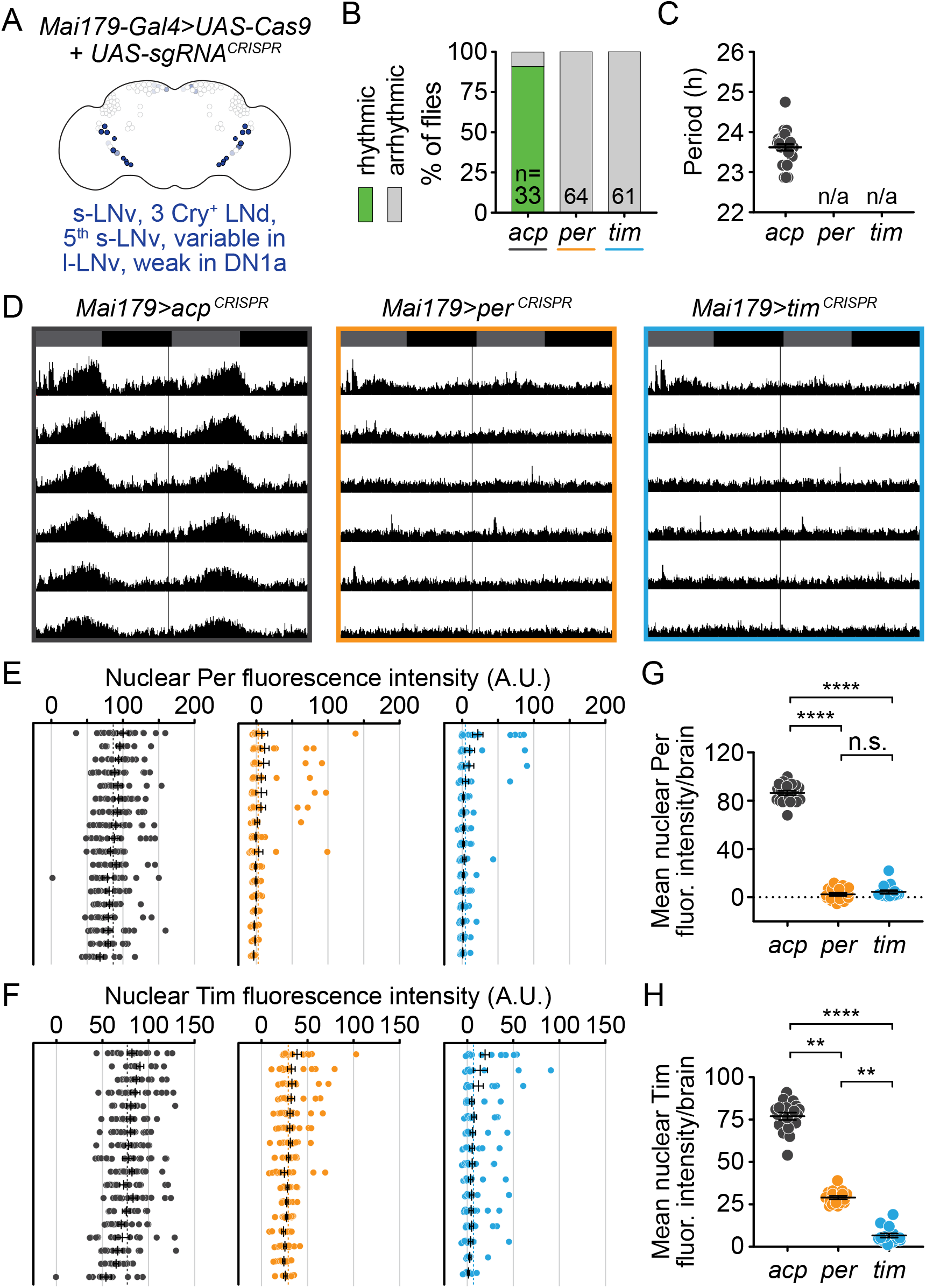
Cell-specific disruption of *per* or *tim* in Mai179^+^ neurons causes complete loss of behavioral rhythmicity and efficient loss of the targeted protein. (A) Diagram of the circadian neuron subset marked by *Mai179-Gal4*. (B) Disruption of *per* or *tim* in Mai179^+^ neurons caused complete loss of behavioral rhythmicity. (C) Scatter plot showing the period of rhythmic flies with *Mai179-Gal4*-driven disruption of *acp*, *per*, or *tim*. (D) Average actograms showing the activity of flies in constant darkness with *Mai179-Gal4*-driven disruption of *acp*, *per*, or *tim*. Six days of activity are displayed. Dark grey rectangles = subjective day, black rectangles = subjective night. (E and F) Background-subtracted nuclear fluorescence intensity of Per (E) or Tim (F) at ZT0 in GFP^+^ neurons, grouped by brain with mean ± SEM shown. (G and H) Mean nuclear fluorescence intensity of Per (G) or Tim (H) at ZT0 in GFP^+^ neurons, averaged per brain (*acp* n = 18, *per* n = 16, *tim* n = 15). ** = p <0.01, **** = p < 0.0001, n.s. = not significant p > 0.05. Significance determined by Kruskal-Wallis nonparametric ANOVA (to account for non-normality of samples) followed by Dunn’s multiple comparisons test; reported p-values are multiplicity adjusted to account for multiple comparisons.

To confirm the loss of protein after gene disruption, we measured Per and Tim protein levels in Mai179^+^ neurons. We co-immunostained brains for Per and Tim and quantified nuclear fluorescence intensity at ZT0. We found that control flies showed robust nuclear staining of both Per and Tim at ZT0 (Fig. 3E-H in grey, S2), whereas *per* disruption in Mai179^+^ neurons caused near-complete loss of Per protein, as shown by the small number of Per^+^ nuclei (Fig. 3E, orange dots) and the average nuclear fluorescence intensity in each brain (Fig. 3G, orange dots). Tim protein in *per* targeted flies remained mostly cytoplasmic (Fig. S2)(17, 46). These results suggest near-complete CRISPR-mediated *per* gene disruption in *Mai179-Gal4*>*per*^*CRISPR*^ flies, consistent with the observed complete loss of behavioral rhythmicity (Fig. 3A-D). For *Mai179*-specific *tim*-targeted flies *(Mai179-Gal4*>*tim*^*CRISPR*^), only a small number of nuclei displayed Tim intensity levels close to the levels observed in control nuclei (Fig. 3F, compare blue to gray), while the average nuclear fluorescence of Tim is near zero (Fig. 3H). Additionally, Per nuclear staining is nearly eliminated in *tim*-targeted flies (Fig. 3E,G). Thus, our results support robust disruption of molecular clock function in these neurons, consistent with the observed complete loss of behavioral rhythmicity.

### Disruption of *per* or *tim* in the morning oscillator (Pdf^+^ s-LNv neurons) does not cause loss of circadian locomotor activity

To investigate the role of the circadian clock in morning oscillator neurons alone, we next induced CRISPR-mediated gene disruption of *tim* and *per* in *Pdf*-expressing cells using *Pdf-Gal4*. Pigment-dispersing factor (Pdf) is a neuropeptide expressed and secreted by the l-LNv and s-LNv neurons, which form the morning oscillator (20, 23, 24, 50) (Fig. 4A). While Pdf^+^ neurons are thought to be essential for circadian locomotor activity, we found that CRISPR-targeting of *per* and *tim* in Pdf^+^ neurons did not cause complete loss of rhythmicity. 74% of *Pdf-*specific, *per*-targeted flies (*Pdf-Gal4*>*per*^*CRISPR*^) and 83% of *Pdf*-specific *tim*-targeted flies (*Pdf-Gal4*>*tim*^*CRISPR*^) were rhythmic, as compared to 100% of controls (*Pdf-Gal4*>*acp*^*CRISPR*^) (Fig. 4B). Qualitatively, activity rhythms of individual flies often appeared more ambiguous, and therefore some were scored as “weakly rhythmic” (Figures 4B, S3). We confirmed these results by an automated scoring method, Lomb-Scargle periodogram analysis (see Methods) (Fig. 4B, S3). Again, *Pdf*-specific targeting of *per* or *tim* resulted in mostly rhythmic flies (87% and 92%, respectively), similar to controls (95% rhythmic) (Table S1). All flies were tracked for 9-10 days after shifting to constant darkness, because *Pdf* mutants and *Pdf receptor* (*Pdfr*) mutants lose their rhythms after 1-3 days in constant darkness (23). We classified flies as rhythmic only if they maintained activity rhythms for the entire 9-10 days. Finally, though the morning oscillator is thought to delay the evening peak of activity and thus control period length (22), the average period length of *Pdf*-specific *per* or *tim*-targeted flies (23.87 and 23.42 hours, respectively) was similar to controls (23.88 hours) (Fig. 4C). These results suggest that the molecular clock is not required in the morning oscillator (Pdf^+^ s-LNv neurons) for overall circadian locomotor activity nor to synchronize other clock neurons such as the evening oscillator.

**Figure 4.**
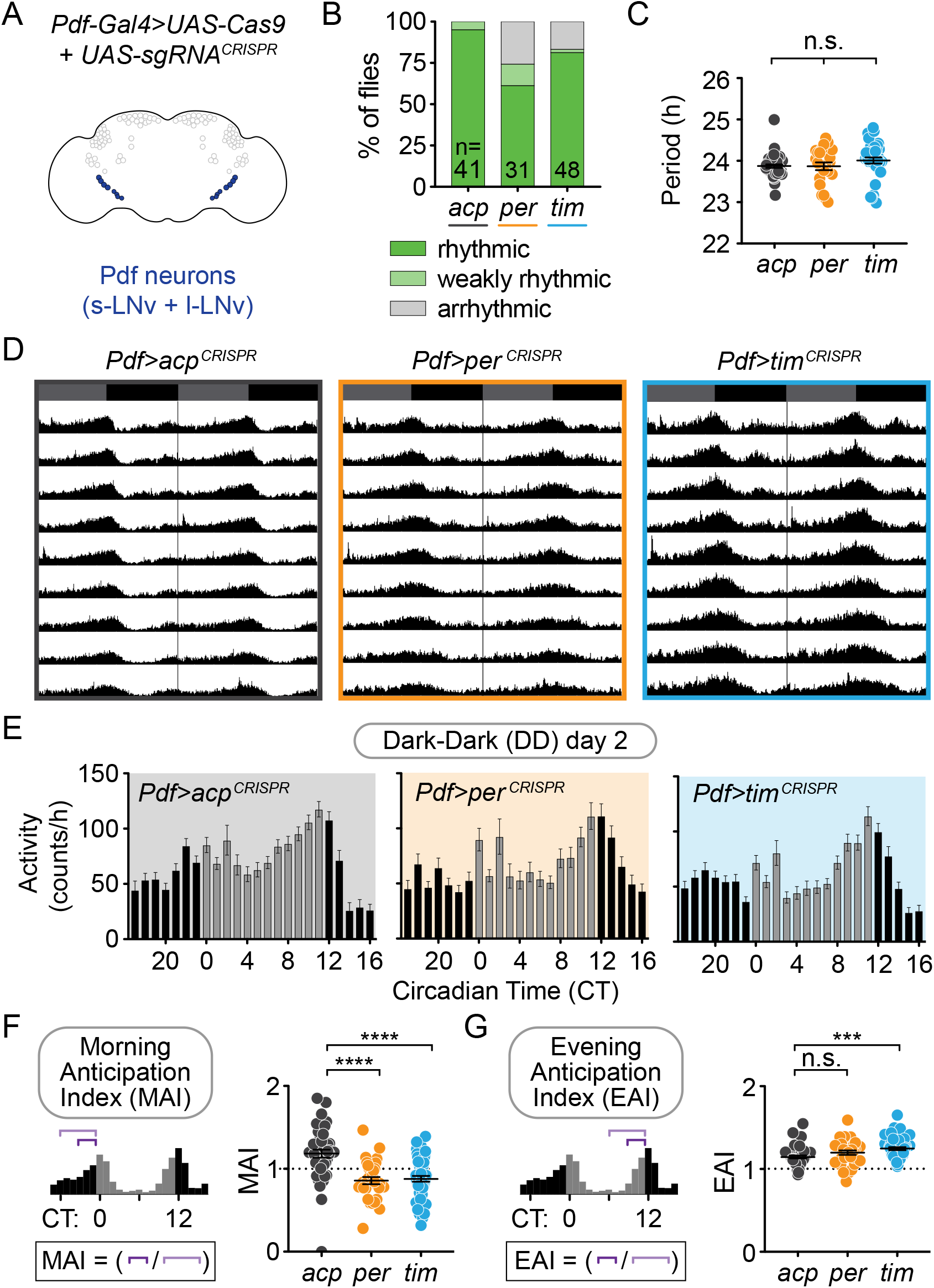
Cell-specific disruption of *per* or *tim* in Pdf^+^ neurons causes incomplete loss of behavioral rhythmicity and loss of morning anticipation in constant darkness. (A) Diagram showing Pdf^+^ circadian neurons. (B) CRISPR-mediated disruption of *per* or *tim* in Pdf^+^ neurons using the *Pdf-Gal4* driver caused incomplete loss of behavioral rhythmicity. (C) Scatter plot showing the period of rhythmic flies with *Pdf-Gal4*-driven disruption of *acp*, *per*, or *tim*. (D) Actograms showing average activity of flies in constant darkness with *Pdf-Gal4*-driven disruption of *acp*, *per*, or *tim*. Nine days of activity are displayed. Dark grey rectangles = subjective day, black rectangles = subjective night. (E) Average hourly activity counts during the second day of complete darkness (DD Day 2; gray bars = CT 0-11, black bars = CT 12-23). Mean number of beam breaks per hour is shown ± SEM. (F) Morning Anticipation Index (MAI) was calculated by dividing the average hourly activity for CT 21-23 by the average hourly activity for CT 18-23. (G) Evening Anticipation Index (EAI) was calculated by dividing the average hourly activity for CT 9-11 by the average hourly activity for CT 6-11. For scatter plots, each point represents an individual fly and mean ± SEM is shown; *** = p < 0.001, **** = p < 0.0001, n.s. = not significant p > 0.05. Significance determined by Kruskal-Wallis nonparametric ANOVA (to account for non-normality of samples) followed by Dunn’s multiple comparisons test; reported p-values are multiplicity adjusted to account for multiple comparisons.

Pdf^+^ s-LNv neurons also regulate “morning anticipation,” or increased activity just before the transition to lights on (20, 21). The evening oscillator neurons regulate “evening anticipation,” or increased activity just before the transition to lights off. To determine whether loss of the molecular clock in Pdf^+^ neurons specifically affects morning anticipation, we analyzed both types of anticipation in *Pdf*-specific *per* and *tim*-targeted flies relative to controls. While evening anticipation was intact after CRISPR-targeting of *tim* or *per* in Pdf^+^ neurons, morning anticipation was absent or diminished (Fig. 4E). To quantify this, we calculated morning and evening anticipation indices (MAIs and EAIs) for each genotype (see Methods). The MAI in DD day 2 was significantly reduced after targeting of *per* or *tim* in Pdf^+^ neurons (*Pdf-Gal4*>*per*^*CRISPR*^ and *Pdf-Gal4*>*tim*^*CRISPR*^) relative to controls (*Pdf-Gal4*>*acp*^*CRISPR*^) (Fig. 4F), whereas EAI was not reduced (Fig. 4G). In LD, the MAI was reduced in *Pdf-Gal4*>*per*^*CRISPR*^ flies, though not in *Pdf-Gal4*>*tim*^*CRISPR*^ flies, and the EAI was unaffected (Fig. S4A,C-F). These phenotypes were similar on days 3-9 in DD (Fig. S4B,G-H). These results suggest that, while the molecular clock in Pdf^+^ neurons is not required for locomotor rhythmicity, it is required for morning anticipatory behavior.

### CRISPR-mediated disruption of *per* or *tim* in Pdf^+^ neurons causes near-complete loss of Per and Tim protein

If the *Pdf-Gal4* driver is not strong enough to fully disrupt *per* or *tim* in Pdf^+^ neurons, this could result in an incomplete behavioral phenotype. To confirm that *per* and *tim* disruption in Pdf^+^ cells is as efficient as observed with *Tim-Gal4* and *Mai179-Gal4*, which caused arrhythmicity, we performed quantitative immunofluorescence analysis of Per and Tim protein levels (Fig. 5). Control flies (*Pdf-Gal4*>*acp*^*CRISPR*^) displayed the expected robust nuclear staining of Per and Tim in both the s-LNvs and l-LNvs (Fig. 5A, D-G, in grey). In contrast, *Pdf-*specific *per*-targeted flies (*Pdf-Gal4*>*per*^*CRISPR*^) exhibited a near-complete absence of nuclear Per immunofluorescence signal in both s-LNvs and l-LNvs (Fig. 4A,D,F). Similar to what we observed in the *Mai179*-specific *per* disruption, any remaining Tim signal was localized to the cytoplasm (Fig 5A). In *Pdf*-specific *tim*-targeted flies (*Pdf-Gal4*>*tim*^*CRISPR*^), we observed a similar near-complete reduction in Tim protein levels in LNvs, with relatively few Tim^+^ nuclei remaining, and an average fluorescence intensity per brain near zero (Fig. 5A,E,G). CRISPR-targeting of *tim* also resulted in near-complete loss of Per protein, indistinguishable from loss of Per in *per*-targeted flies (Fig. 5F)(46, 47). These results suggest that the persistence of circadian locomotor activity seen in *Pdf*-specific *per* and *tim*-targeted flies is not due to incomplete disruption of the targeted gene. Taken together, our behavioral and quantitative immunofluorescence analysis suggest that the molecular clock in Pdf^+^ clock neurons is not required for circadian locomotor activity.

**Figure 5.**
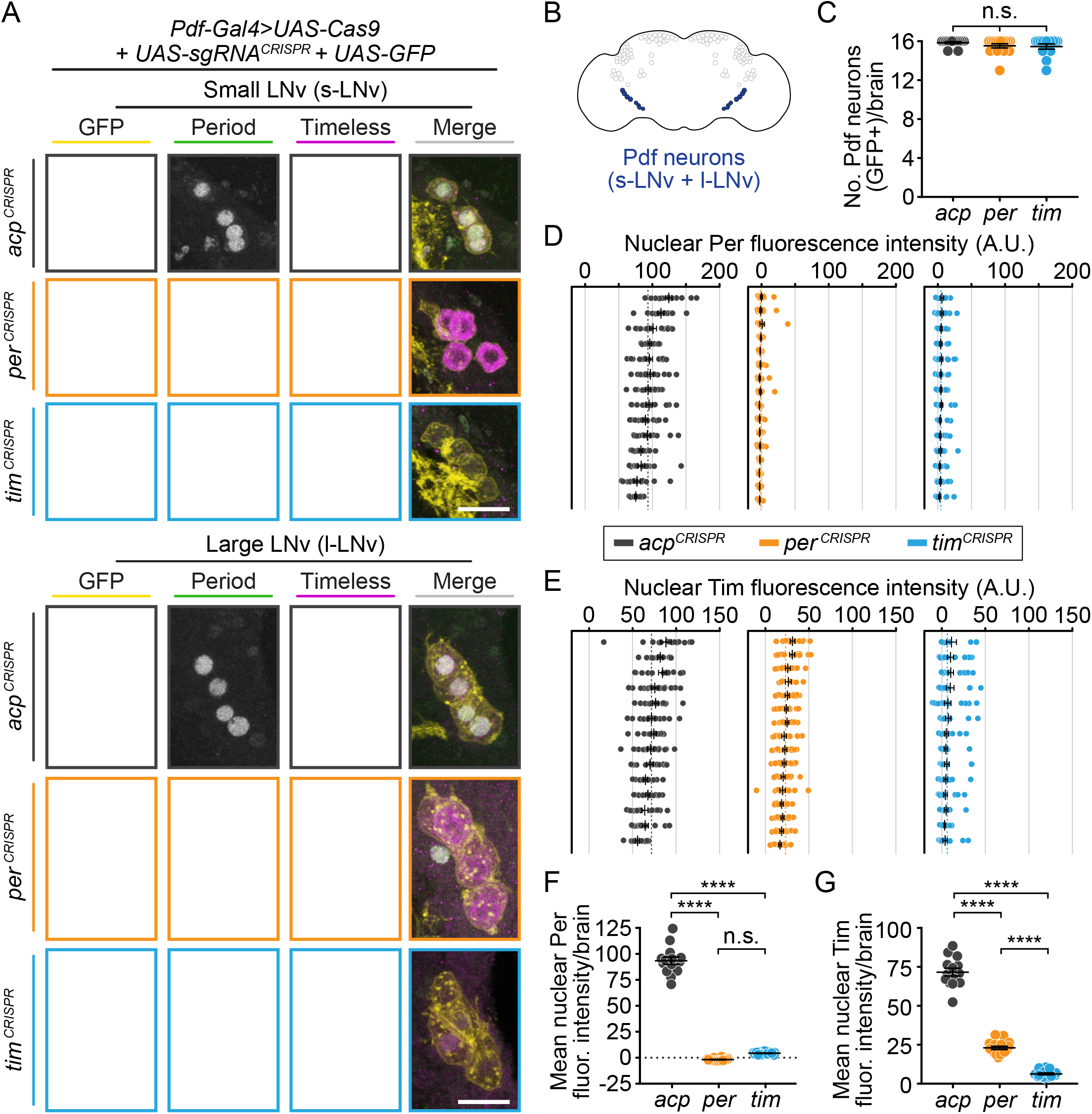
CRISPR-mediated disruption of *per* or *tim* in Pdf^+^ neurons leads to efficient loss of the targeted protein. (A) Maximum intensity projections of *Pdf-Gal4*-driven GFP^+^ small and large ventral lateral neurons (s- and l-LNv) with immunohistochemistry for GFP (yellow), Period (green) and Timeless (purple) at ZT0. Scale bar = 10 μm; arrows indicate CRISPR-targeted nuclei with residual protein signal. (B) Diagram showing *Pdf-Gal4* expression in the 4 large and 4 small ventral lateral neurons (l- and s-LNv), bilaterally. (C) Quantification of the number of GFP^+^ neurons per brain (*acp* n = 14, *per* n = 15, *tim* n = 13; brains scored only if both hemispheres were intact after dissection). (D and E) Background-subtracted nuclear fluorescence intensity of Per (D) or Tim (E) at ZT0 in GFP^+^ neurons, grouped by brain with mean ± SEM shown. (F and G) Mean nuclear fluorescence intensity of Per (G) or Tim (H) at ZT0 in GFP^+^ neurons, averaged per brain (*acp* n = 14, *per* n = 16, *tim* n = 14). **** = p < 0.0001, n.s. = not significant p > 0.05. Significance was determined by one-way ANOVA followed by Tukey’s multiple comparisons test; reported p-values are multiplicity adjusted to account for multiple comparisons.

We also used a *UAS-myr-GFP* to label the membranes of Pdf^+^ neurons and counted the number of GFP^+^ cells in each brain to confirm that the CRISPR-induced DNA damage in our system does not cause cell death. There are 8 Pdf^+^ LNvs in each hemisphere, totaling 16 neurons in each fly brain (51). We found no significant difference in the number of GFP^+^ LNvs in each brain between experimental flies and controls (Fig. 5C). This result indicates that CRISPR-mediated gene disruption in Pdf^+^ neurons does not cause significant cell death.

### Restriction of CRISPR-mediated disruption of *per* or *tim* to evening oscillator (Mai179^+^ Pdf^−^) neurons by blocking disruption in Pdf^+^ neurons restores behavioral rhythmicity

To determine whether the molecular clock is required in evening oscillator neurons, we paired the *Mai179-Gal4* driver with *Pdf-Gal80*, blocking CRISPR-targeting of *tim* or *per* in Pdf^+^ neurons. This restricts CRISPR-targeting to the evening oscillator (Mai^+^ Pdf^−^ neurons): the 5^th^ sLNv, CRY^+^ LNds, and a small subset of DN1s (Fig. 6A). We found that overall locomotor rhythmicity is maintained; 86% of *per*-targeted flies (*Mai179-Gal4/Pdf-Gal80*>*per*^*CRISPR*^) and 85% of *tim*-targeted flies (*Mai179-Gal4/Pdf-Gal80*>*tim*^*CRISPR*^) were rhythmic, similar to 91% of control flies (Fig. 6B-D). This result is consistent with previous studies in which rhythmicity is restored by expression of *UAS-per* in Pdf^+^ neurons of *per* null mutant flies (21). Thus, while *per* and *tim* expression are not necessary in Pdf^+^ neurons for rhythmicity (Fig. 4), their expression in Pdf^+^ neurons is sufficient for circadian locomotor activity. Taken together, our results suggest that the molecular clock is sufficient in either the morning oscillator or evening oscillator for circadian locomotor activity and that the molecular clock must be disrupted in both oscillators to disrupt circadian locomotor activity.

**Figure 6.**
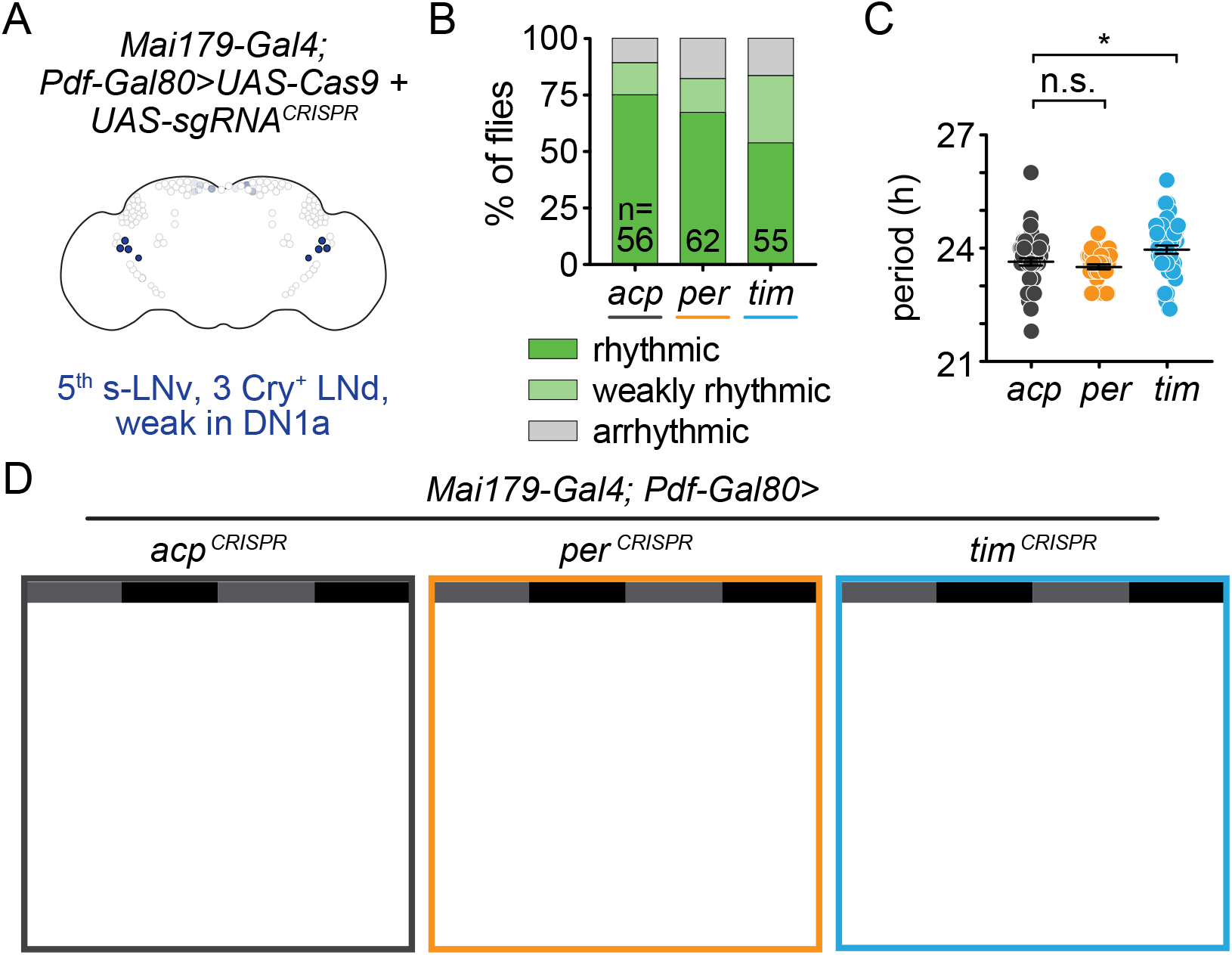
Cell-specific disruption of *per* or *tim* in Mai179^+^Pdf^−^ neurons does not affect behavioral rhythmicity. (A) Diagram showing circadian neurons targeted using *Mai179-Gal4; Pdf-Gal80*. (B) CRISPR-mediated disruption of *per* or *tim* in Mai179^+^Pdf^−^ neurons did not affect behavioral rhythmicity. (C) Scatter plot showing the period of rhythmic flies with *Mai179-Gal4; Pdf-Gal80*-driven disruption of *acp*, *per*, or *tim*. (D) Average actograms showing the activity of flies in constant darkness with *Mai179-Gal4; Pdf-Gal80*-driven disruption of *acp*, *per*, or *tim*. Six days of activity are displayed. Dark grey rectangles = subjective day, black rectangles = subjective night.

Because Mai179^+^ Pdf^−^ neurons comprise the minimal evening oscillator, we also measured “evening anticipation,” or increased activity just before the transition to lights off. Similar to our observation that *per* or *tim* disruption in the Pdf^+^ morning oscillator led to a loss of morning anticipation, *per* or *tim* disruption in the Mai^+^ Pdf^−^ evening oscillator neurons led to a loss of evening anticipation activity (Fig. 6E,G). The EAI was significantly reduced in evening oscillator-specific *per*-targeted flies and the EAI of *tim*-targeted flies was trending, but not significantly reduced (p<0.10), relative to controls. The morning anticipation indices (MAI) remained intact and were not significantly different from controls (Fig. 6F), further demonstrating that *per* or *tim* expression in Pdf^+^ s-LNv morning oscillator neurons is both sufficient and necessary for morning anticipatory activity.

## Discussion

Our understanding of how circadian neurons communicate with each other to control locomotor rhythmicity is still evolving. Over a decade ago, some of the first evidence was presented to support a “dual oscillator” model in which Pdf^+^ s-LNvs are classified as “morning cells” that control morning anticipation and drive the maintenance of overall rhythmicity. This model also suggests that Pdf^+^ s-LNvs dominate over other circadian neurons, such as “evening cells” (19–21, 41). More recent evidence has questioned this hierarchical model and instead suggests a complex network in which the control of circadian behavior is distributed among many subgroups of neurons (22, 27, 52–54). Most of these recent studies utilized proteins known to alter period length when expressed ubiquitously (mutant kinases, mutated kinase targets, or dominant negative constructs). When these proteins were overexpressed in specific clock neurons such as Pdf^+^ neurons or evening oscillators, they exerted varying levels of control over the molecular clocks in other neurons and overall circadian locomotor activity. While these studies were elegantly done using available tools, overexpression studies carry the potential problem of gain of function. Moreover, proteins that regulate the core molecular clock have significantly different roles in different clock neurons (55, 56). The best genetic tools are those that cause loss of function. Here we developed and validated tools for CRISPR-mediated disruption of the molecular clock in targeted subsets of circadian neurons.

Our results demonstrate the efficacy and utility of genetic constructs that mediate tissue-specific CRISPR-targeting of two key circadian clock genes: *timeless* and *period*. We showed that these constructs recapitulate known mutant phenotypes, such as complete loss of locomotor activity rhythms when driven in all *tim*-expressing cells (Fig. 2). We validated the extent of gene disruption at both the mRNA and protein levels (Fig. 2, 3, and 5) and showed that gene targeting effects are cell-type specific, as there is no effect on locomotor rhythmicity when *tim* or *per* are disrupted in glia. We then used these lines to dissect the molecular clock requirements in different subsets of circadian regulatory neurons. Our results show that *per* and *tim* expression is sufficient but not necessary for circadian locomotor activity in either Pdf^+^ cells (which include the morning oscillator) or the evening oscillator neurons (Mai^+^ Pdf^−^), though loss of *per* or *tim* expression in both Pdf^+^ neurons and the evening oscillator leads to arrhythmicity.

While this manuscript was in preparation, we became aware of a similar study examining the requirement of the molecular clock in different subsets of clock neurons. Schlichting *et al.* also used a tissue-specific CRISPR-mediated mutagenesis strategy to target *period* with three gRNAs and obtained similar results. Consistent with our results, they found that disruption of *per* expression in Pdf^+^ cells did not cause loss of circadian locomotor activity. Moreover, loss of Clock protein cycling in Pdf^+^ neurons due to Pdf-specific neuronal silencing also did not cause loss of circadian locomotor activity. Taken together, our results and those from the Rosbash lab demonstrate that the molecular clock is not required in Pdf^+^ neurons for circadian locomotor activity and suggests that rhythmicity is a network property.

Our evidence supports a model of independent morning and evening oscillators that control their respective anticipatory behaviors, but can compensate for each other to maintain overall locomotor rhythmicity. CRISPR targeting *per* or *tim* in Pdf^+^ neurons, which contain the morning oscillator, led to a loss of morning anticipatory behavior (Fig. 4). This is consistent with previous reports demonstrating that ablation of *Pdf*-expressing cells or loss of function of *Pdf* itself or its receptor *Pdfr* caused loss of morning anticipation (23). This suggests that this specific aspect of circadian behavior, morning anticipatory activity, requires the molecular clock in Pdf^+^ neurons. However, while *Pdf* mutants and Pdf^+^ cell ablation led to a loss of overall rhythmicity (23, 25, 29, 40), disruption of the molecular clock (*tim* or *per*) in Pdf^+^ neurons did not. These results suggest that locomotor rhythmicity, while dependent on *Pdf* expression and Pdf^+^ neurons, is not dependent on the function of the molecular clock within Pdf^+^ neurons. Similarly, disruption of *per* or *tim* in just the evening oscillator neurons (*Mai179-Gal4/Pdf-Gal80*), led to a loss of evening anticipatory behavior, but not locomotor rhythmicity (Fig. 6). This also demonstrated that while an intact molecular clock in morning oscillator neurons was not necessary for overall rhythmicity, it was sufficient to restore the rhythmicity lost with *Mai179-Gal4-*driven disruption of *per* or *tim*. These results are consistent with recent work suggesting that interactions between clock neurons create multiple independent oscillators that regulate locomotor activity rhythms (27).

Our results further suggest that the molecular clock needs to be disrupted in both the morning and evening oscillator neurons to disrupt locomotor rhythmicity. When we drove *per*^*CRISPR*^ or *tim*^*CRISPR*^ with *Mai179-Gal4*, which expresses in a subset of clock neurons that include both morning oscillator neurons (s-LNvs) and evening oscillator neurons (primarily 3 CRY^+^ LNds, and the 5^th^ s-LNv), we saw a complete loss of overall rhythmicity. Previous research has shown that rescuing the circadian clock with *UAS-per* expression in a *per* null background in *Mai179-Gal4* cells was not sufficient to fully restore rhythmicity (21, 50, 57), but it is possible that UAS-driven expression of *per* did not fully recapitulate endogenous, cyclical expression levels. In contrast, our results demonstrate that the molecular clock in one or more of the *Mai179-Gal4* expressing neurons is necessary for behavioral rhythms. Perhaps the morning and evening oscillators function with some redundancy, coordinating rhythmicity in a distributed, complex network, that only requires a cell-intrinsic molecular clock in one subset of neurons to generate behavioral rhythms. In other words, an intact molecular clock in one subset of clock neurons is able to compensate for loss in another subset, suggesting that clock neurons do not rely entirely on a cell-intrinsic molecular clock to generate behavioral rhythms.

Our results highlight how cell-specific CRISPR-mediated gene disruption can be used to better understand the role the molecular clock plays in specific subsets of circadian neurons to control behavioral rhythmicity. Our work also demonstrates the immense potential of the approach engineered by Port & Bullock to produce cell-specific, CRISPR-mediated gene disruption in somatic cells. These tools provide a new standard for the field and can now be used to investigate the tissue-specific function of circadian genes in both neuronal subsets and “peripheral clocks” outside the brain that control other circadian-regulated physiologies.

## Author contributions

RD, MP, RMO, and MSH conceived of and designed experiments. RD, MP, RMO, MU and MB performed experiments. RD, MP, RMO, MU and MB analyzed and interpreted data. HXK generated tools and provided technical support. RD, MP, RMO, JCC, and MSH prepared manuscript.

## Acknowledgements

This work was supported by the following funding sources: NIH R01GM105775 (M.S.H.), R35 GM127049 (M.S.H.), NIH R01GM117407 (J.C.C.), NIH 2T32GM007367-42 (R.M.O., MSTP training grant), 5T32HL120826 (R.D.), NIH training grant 5T32DK007328 (M.U.), and the Charles H. Revson Foundation Senior Postdoctoral Fellowship in Biomedical Science (M.U.).

Stocks obtained from the Bloomington *Drosophila* Stock Center (NIH P40OD018537) were used in this study.

## Materials & Methods

### Drosophila *strains and maintenance*

*UAS-sgRNA* lines (*w;UAS-sgRNA-tim*^*3x*^; *w;UAS-sgRNA-per*^*4x*^; and *w;UAS-sgRNA-acp98AB*^*4x*^;) were cloned as described below. The *w;;UAS-Cas9.2* line was obtained from Bloomington *Drosophila* Stock Center (#58986). Two different *UAS-myr-GFP* lines were used (2^nd^ chromosome: Bloomington #32198 and 3^rd^ chromosome: Bloomington #32197). *per*^01^ nulls were a gift from Jaga Giebultowicz.

*Gal4* drivers: *w;tim-Gal4;* (Bloomington #7126), *w;;repo-Gal4* (Bloomington #7415), w;*Mai179-Gal4*; (Helfrich-Förster Lab), w;*Pdf-Gal4;* (Bloomington #6900). All driver lines were outcrossed at least six generations to *w*^*−*^*CS* (white-eyed *Canton-S* strain).

All flies were grown and maintained on standard yeast-cornmeal-agar media (Archon Scientific, Glucose recipe: 7.6% w/v glucose, 3.8% w/v yeast, 5.3% w/v cornmeal, w/v 0.6% agar, 0.5% v/v propionic acid, 0.1% w/v methyl paraben, 0.3% v/v ethanol) in a humidity controlled (55-65%) 12:12 Light:Dark incubator at 25°C. Males were collected at 1-3 days old and allowed to mate for 1-2 days before being separated from females. Male flies were 7-11 days old at the start of all behavioral and immunohistochemistry experiments.

### Cloning

Multiple gRNAs targeting *per*, *tim*, or *acp98AB* were constructed as previously described (35). gRNA sequences were selected for predicted target specificity and efficiency according to http://chopchop.cbu.uib.no/ (58). *pCFD6* (Addgene #73915) was digested with BbsI-HF (NEB #R3539S) and gel purified. For each construct, inserts were generated in three separate PCR reactions using *pCFD6* as the template and the primers listed in Table S2. The resulting three inserts and the pCFD6 backbone were then assembled by NEBuilder HiFi DNA Assembly (NEB #E2621L) for each construct. Each construct was integrated at the *Su(Hw)attP5* site (59) (Bestgene, Inc.) and Sanger sequenced (Genewiz). Sequenced flies revealed a polymorphism in one of the four sgRNA scaffolds in the *UAS-t:sgRNA-tim* flies and thus the line is denoted as *UAS-t:sgRNA-tim*^*3x*^.

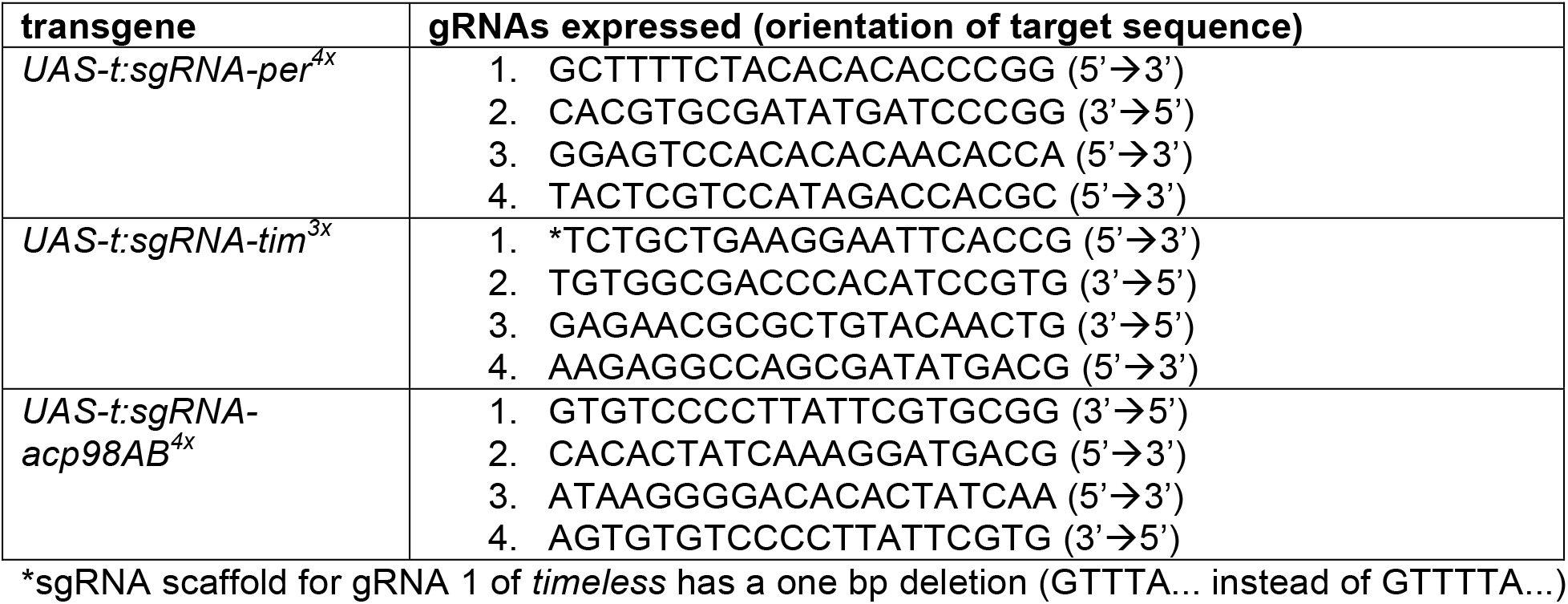

### Circadian locomotor activity

Male flies entrained on a 12:12 LD cycle during development and post-eclosion were placed in individual 5mm tubes in TriKinetics, Inc. *Drosophila* Activity Monitors (DAMs) to record their locomotor activity for two days in 12:12 LD, then for 7-11 days in constant darkness (DD). Activity data from the DD period was grouped into 15 min bins and Clocklab software (Actimetrics) was used to generate actograms and period measurements. Actograms were blindly scored as rhythmic, weakly rhythmic, or arrhythmic; percentages of each category are reported, except when “weakly rhythmic” was less than 5% of the population, then it was included with “rhythmic.” To calculate Anticipatory Indices (MAI or EAI), activity for individual flies was binned into 1-hour windows for each individual fly. An anticipatory index was calculated by dividing the average activity per hour over 3 hours immediately preceding “lights on” (MAI) or “lights off” (EAI) by the average activity per hour over the 6 hours preceding “lights on” (MAI) or “lights off” (EAI), in LD day 2, DD day2, and DD days 3-9. All circadian data is represented as a sum of at least three biological replicates of at least 8-10 flies each per genotype.

### Automated Circadian Analysis

Using the Clocklab Actimetrics Software, Lomb-Scargle periodograms with a statistical cutoff of p<0.001 were generated. The difference between the amplitude of the peak and the value of the best fit line was calculated and flies were classified as rhythmic if the difference was >150. The % rhythmicity results are reported in Table S1.

### Immunohistochemistry and Confocal Microscopy

After 6-9 days of entrainment, flies were decapitated at ZT0 and heads were fixed in 4% paraformaldehyde (Electron Microscopy Sciences #RT15710) in PBS + 0.1% Triton X-100 (PTX) for 40 minutes at room temperature. Heads were washed in PTX and subsequently incubated on ice. Brains were dissected in PTX and blocked with 4% normal donkey serum (NDS, Jackson ImmunoResearch #017-000-121) in PTX for 90 minutes at room temperature. After blocking, brains were incubated overnight at 4°C in primary antibody: chicken α-GFP (1:1000, Abcam #ab13970), rabbit α-Per (1:1000, gift of Michael Rosbash (60)), and rat α-Tim (1:1000, gift of Amita Sehgal and Michael Young (13)) in PTX + 2% NDS. Rabbit α-Per was pre-adsorbed on dechorionated *per*^*01*^ embryos overnight in PTX + 2% NDS prior to use. Brains were washed in PTX and incubated overnight at 4°C in secondary antibody: Alexa Fluor 488–conjugated donkey α-chicken (1:200, Jackson ImmunoResearch #703-545-155), Alexa Fluor 594–conjugated donkey α-rabbit (1:200, Jackson ImmunoResearch #711-585-152), and Alexa Fluor 647–conjugated donkey α-rat (1:200, Jackson ImmunoResearch #712-605-153) in PTX + 2% NDS. Brains were washed in PTX then PBS and were mounted on coverslips coated with Poly-L-Lysine (0.1 mg/mL, Advanced BioMatrix #5048) and PhotoFlow 200 (0.36%, Kodak #1464510). Coverslips were serially dehydrated with increasing concentrations of ethanol (30, 50, 75, 95, 100, 100%) and cleared with two washes in 100% xylenes. Coverslips were mounted onto slides with DPX (Electron Microscopy Sciences #RT13510) and dried overnight before imaging.

Images were acquired on a Zeiss LSM 800 Axio Observer 7 inverted confocal microscope (ZEISS) using 488-, 561-, and 647-nm lasers and a Plan-Apochromat 63x/1.40 oil immersion lens (Fig. 3, 5, S2) or 20x/0.8 dry lens (Fig. S1). Z-stacks were taken using Zeiss LSM confocal software Zen 2.3 (1.5 μm slice thickness). Image analysis was performed in FIJI (61); mean fluorescence intensity of GFP positive nuclei was measured, normalized by subtracting a measurement of mean background intensity, and analyzed using GraphPad Prism software. For *Pdf-Gal4* experiments, the number of GFP^+^ neurons in each brain was counted and analyzed to assess potential CRISPR-driven cytotoxicity.

### Quantitative real-time PCR (QRT-PCR)

14-day old male flies previously entrained to 12:12 LD were placed in constant darkness (DD) for 24 hours, after which 7 circadian timepoints were taken at CT-1, 5, 9, 13, 17, 21 and 25, and snap frozen in liquid nitrogen and stored at −80°C. RNA was extracted from 60 heads for each of 4 biological replicates per genotype/timepoint with TRIzol (Invitrogen) following the manufacturer’s protocol. Samples were treated with DNaseI (Invitrogen) then heat inactivated. cDNA was synthesized by Revertaid First Strand cDNA Synthesis Kit (Thermo Scientific). PowerUp SYBR Mastermix (Applied Biosystems) was used to perform QRT-PCR using a CFX-Connect thermal cycler (BioRad). Primer efficiency and relative quantification of transcripts was determined using a standard curve of serial diluted cDNA. Transcripts were normalized using Actin5C as a reference gene.

Primer sequences:

*clock*-fwd-GGATAAGTCCACGGTCCTGA
*clock*-rev-CTCCAGCATGAGGTGAGTGT
*period*-fwd-CGAGTCCACGGAGTCCACACACAACA
*period*-rev-AGGGTCTGCGCCTGCCC
*timeless*-fwd-CCGTGGACGTGATGTACCGCAC
*timeless*-rev-CGCAATGGGCATGCGTCTCTG
*Actin5C*-fwd-TTGTCTGGGCAAGAGGATCAG
*Actin5C-*rev-ACCACTCGCACTTGCACTTTC

## Supplementary Figure Legends

**Supplementary Figure 1.**
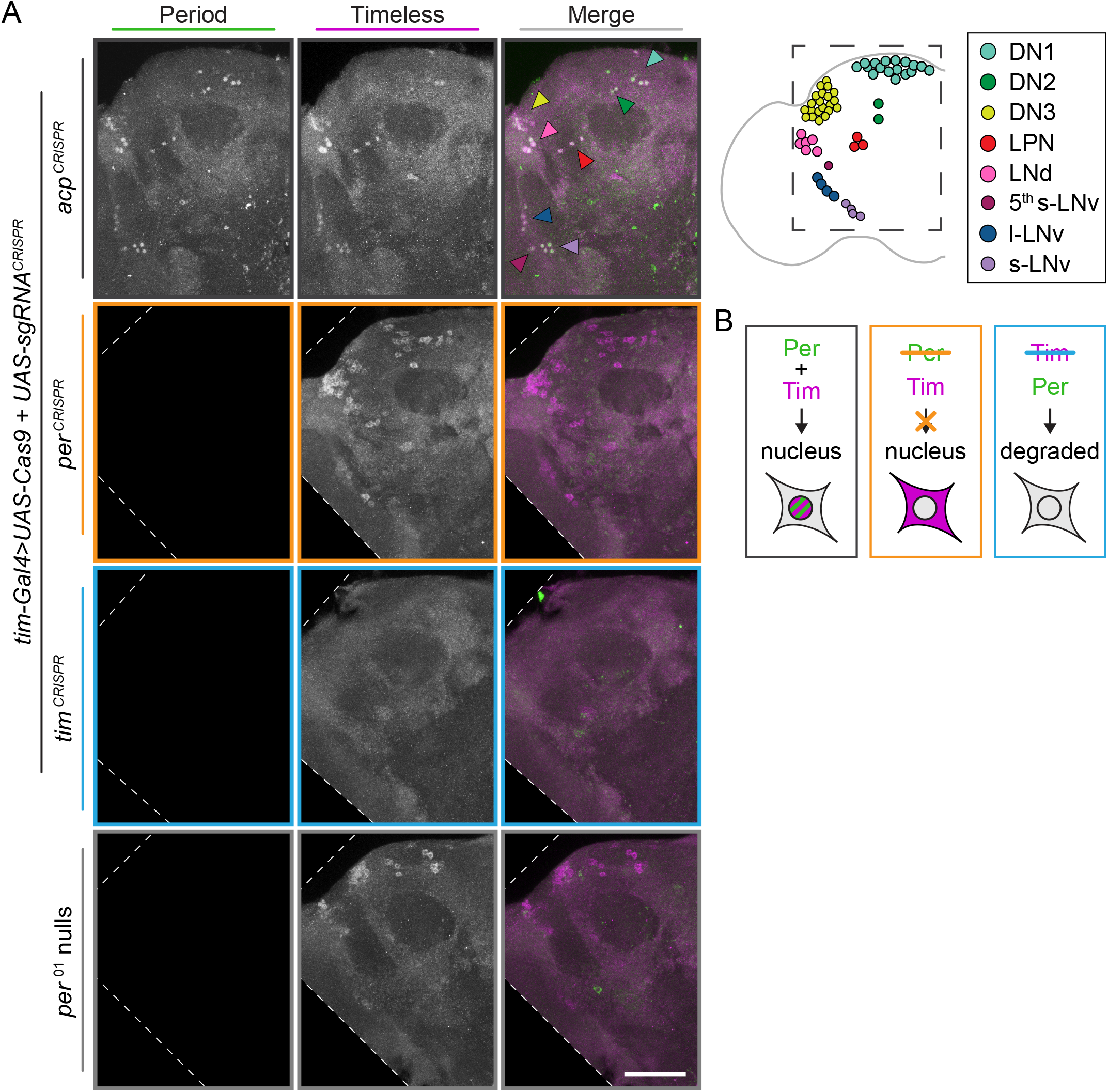
CRISPR-targeting of *per* or *tim* in Tim^+^ neurons leads to overall loss of Per and Tim protein. (A) Maximum intensity projections of *tim-Gal4*>*acp*^*CRISPR*^, *tim-Gal4*>*per*^*CRISPR*^, *tim-Gal4*>*tim*^*CRISPR*^, and *per*^01^ null brains at ZT0 with immunohistochemistry for Period (green) and Timeless (purple) with schematic of clock neuron clusters; scale bar = 50 μm. (B) Schematic showing expected staining pattern based on Per and Tim stability and requirements for nuclear entry.

**Supplementary Figure 2.**
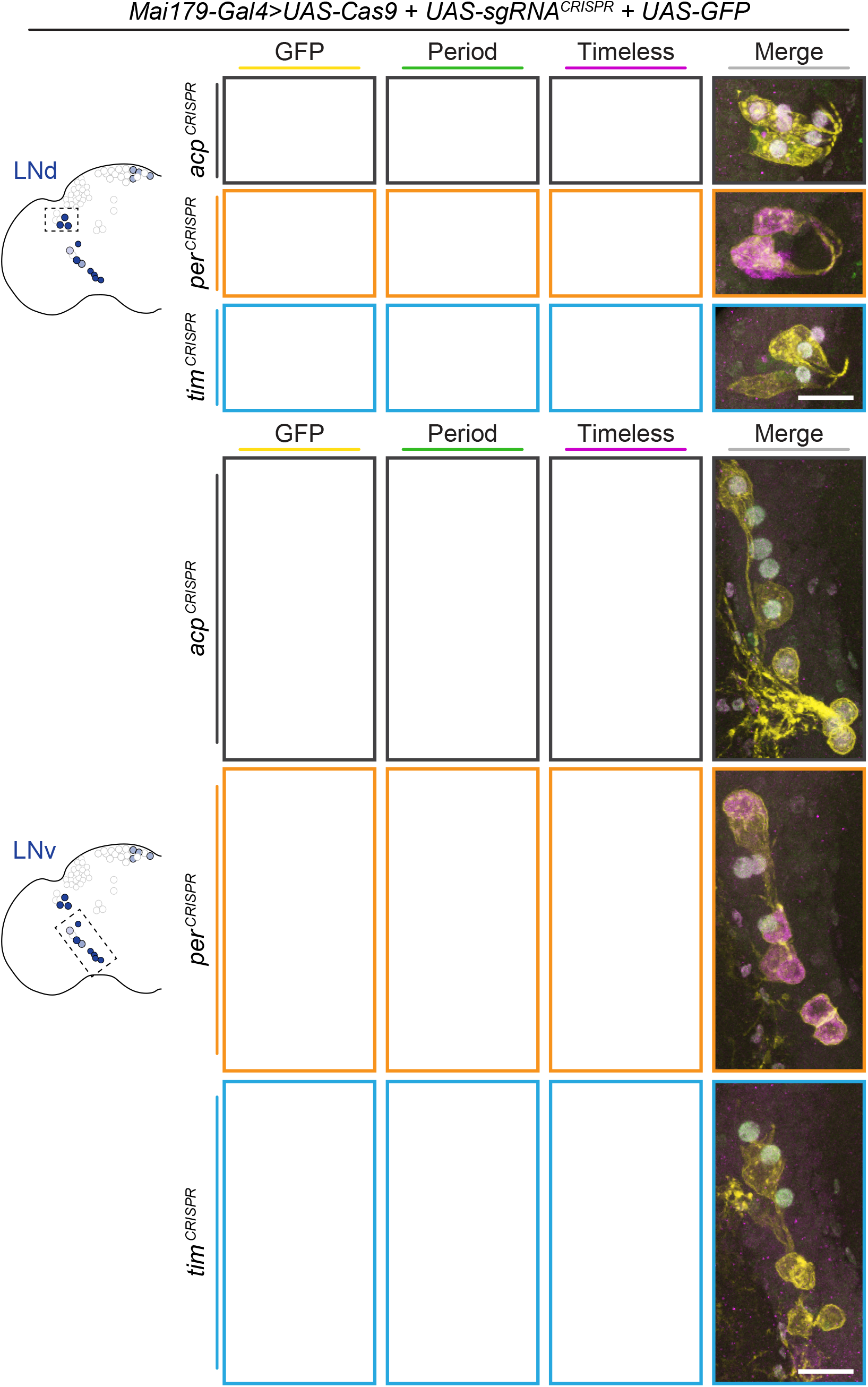
Period and Timeless immunohistochemistry in *Mai179-Gal4*-driven CRISPR flies. Maximum intensity projections of *Mai179-Gal4*-driven GFP^+^ dorsal lateral (LNd) and ventral lateral (LNv) neurons with immunohistochemistry for GFP (yellow), Period (green) and Timeless (purple); scale bar = 10 μm.

**Supplementary Figure 3.**
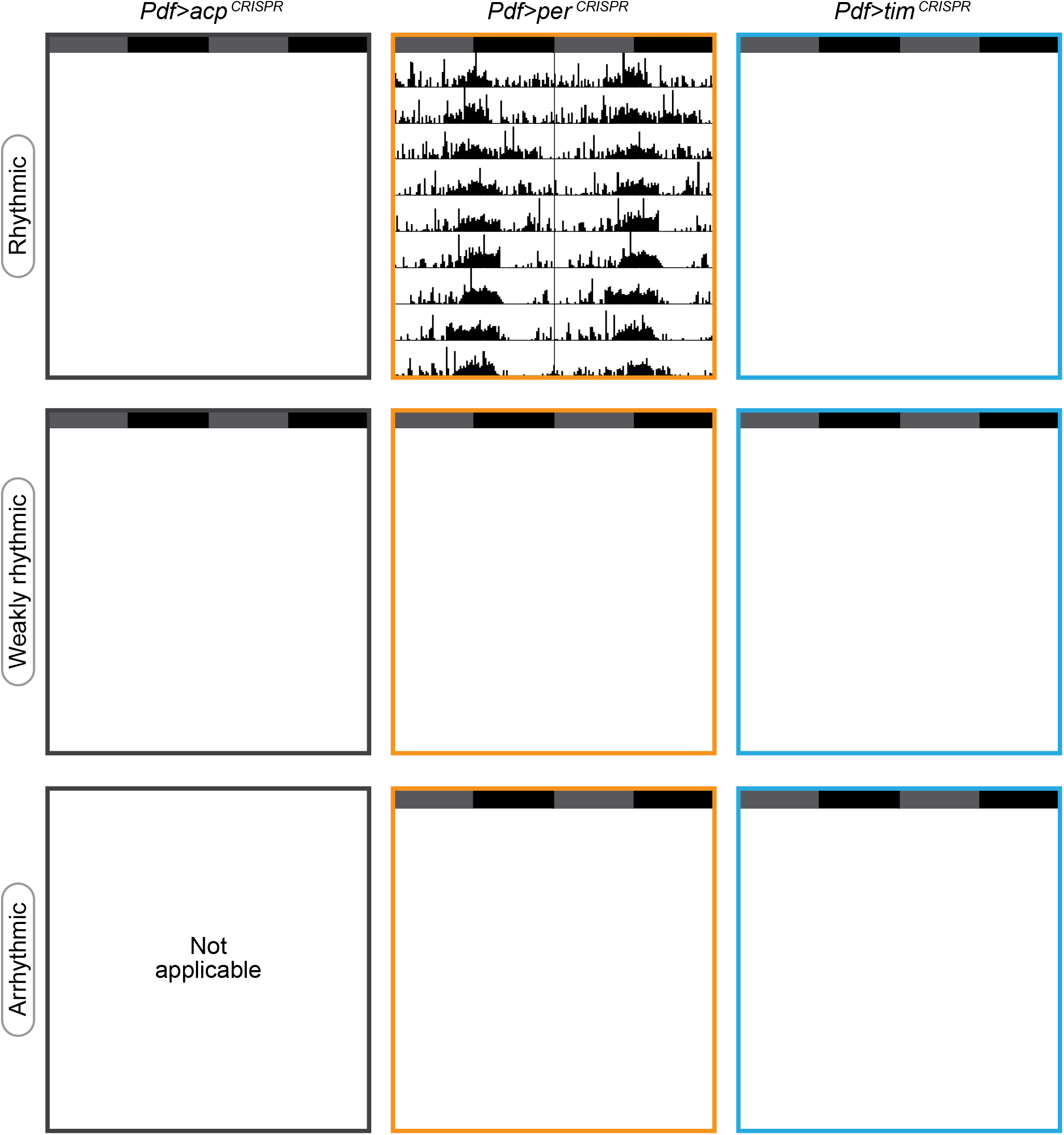
Representative actograms for each phenotypic class. Representative single-fly actograms of *Pdf-Gal4*>*acp*^*CRISPR*^ (gray), *per*^*CRISPR*^ (orange), or *tim*^*CRISPR*^ (blue) flies blindly scored as rhythmic (top row), weakly rhythmic (middle row), or arrhythmic (bottom row; no *Pdf-Gal4*>*acp*^*CRISPR*^ flies were scored as arrhythmic).

**Supplementary Figure 4.**
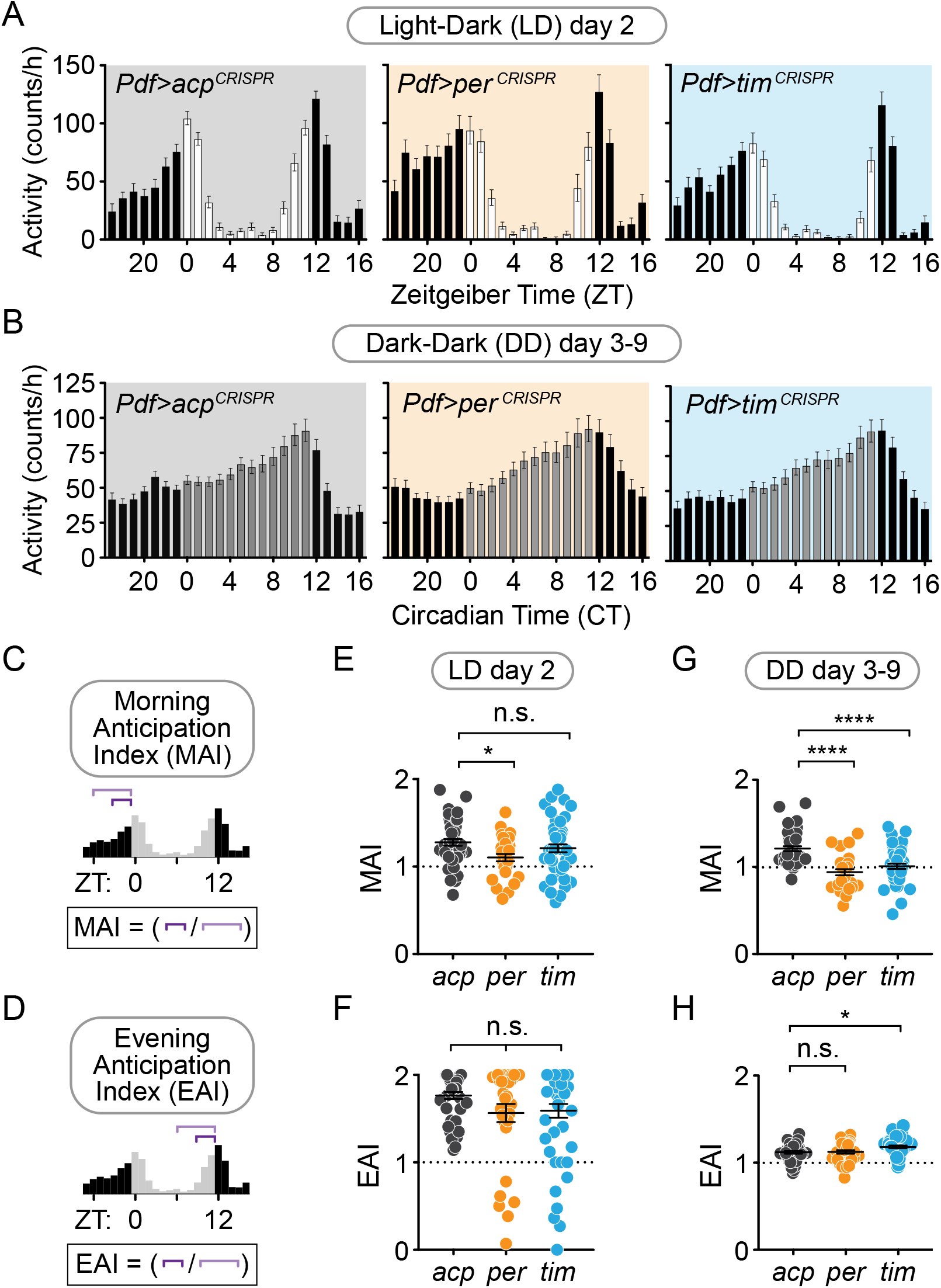
Cell-specific disruption of *per* or *tim* in Pdf^+^ neurons causes loss of the morning anticipatory peak under constant conditions. (A and B) Average hourly activity counts for one day under a 12h:12h light-dark schedule (A; white bars = ZT 0-11, black bars = ZT 12-23) or averaged over days 3-9 of complete darkness (B; gray bars = CT 0-11, black bars = CT 12-23); mean number of beam breaks per hour is shown ± SEM. (C) Morning Anticipation Index (MAI) calculated by dividing the average hourly activity for ZT (E) or CT (G) 21-23 by the average hourly activity for ZT or CT 18-23. (*acp* n = 42, *per* n = 31, *tim n* = 48). (D) Evening Anticipation Index (EAI) calculated by dividing the average hourly activity for ZT (F) or CT (H) 9-11 by the average hourly activity for ZT or CT 6-11. (E and F) Morning (E) and Evening (F) Anticipation Indices for one day under a 12h:12h light-dark schedule. (*acp* n = 42, *per* n = 31, *tim n* = 48). (G and H) Average Morning (G) and Evening (H) Anticipation Indices over days 3-9 under constant darkness. (*acp* n = 42, *per* n = 31, *tim n* = 48). For scatter plots, each point represents an individual fly and mean ± SEM is shown. Significance determined by one-way ANOVA followed by Tukey’s multiple comparisons test (for normally distributed samples; E and H) or Kruskal-Wallis nonparametric ANOVA (to account for non-normality of samples; F and G) followed by Dunn’s multiple comparisons test; reported p-values are multiplicity adjusted to account for multiple comparisons. * = p < 0.05, **** = p < 0.0001, n.s. = not significant p > 0.05.

**Supplementary Figure 5.**
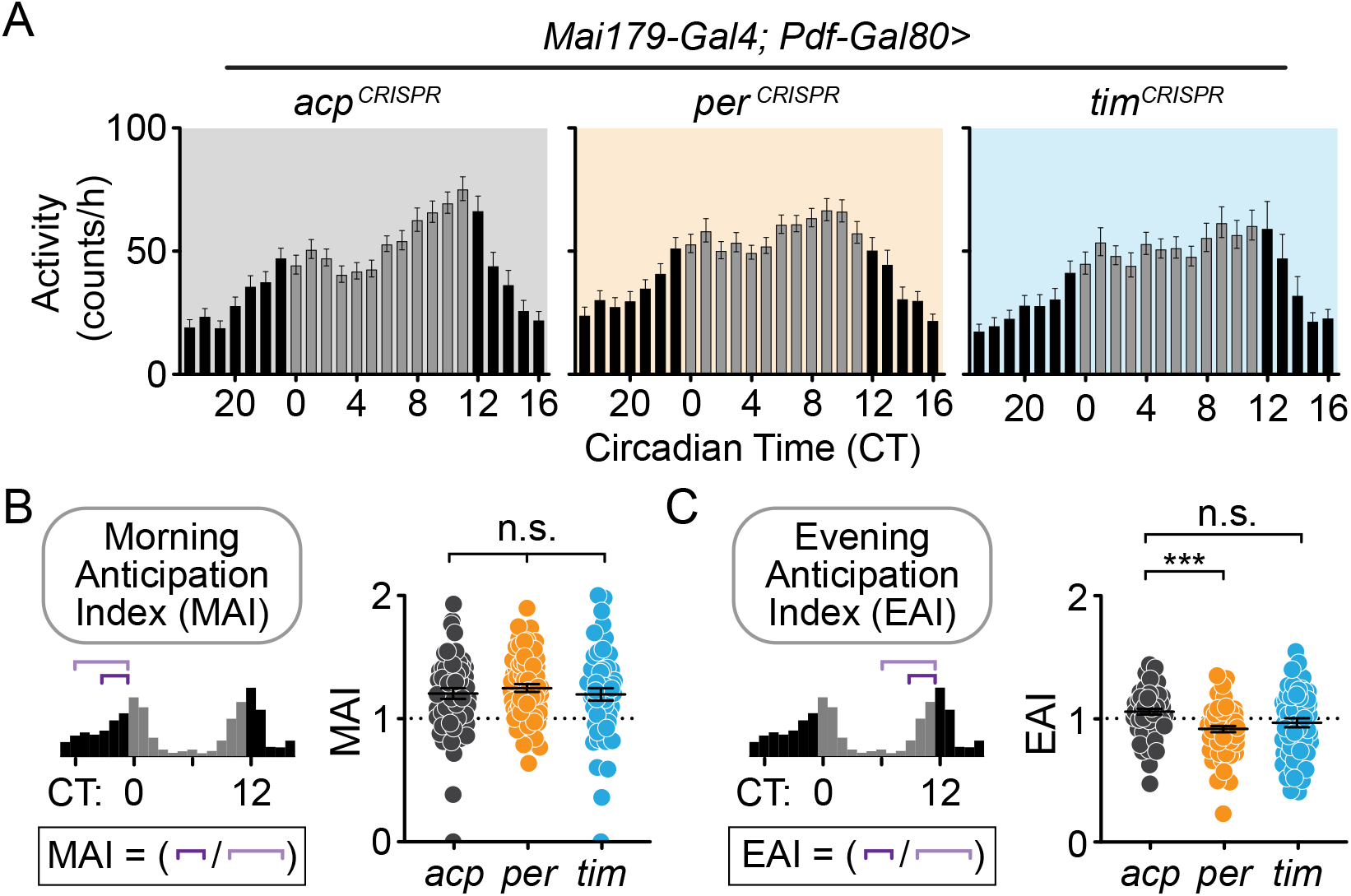
Cell-specific disruption of *per* in Mai179^+^Pdf^−^ neurons causes loss of the evening anticipatory peak under constant conditions. (A) Average hourly activity counts during the second day of complete darkness (DD Day 2; gray bars = CT 0-11, black bars = CT 12-23). Mean number of beam breaks per hour is shown ± SEM. (*acp* n = 58, *per* n = 63, *tim n* = 55) (B) Morning Anticipation Index (MAI) was calculated by dividing the average hourly activity for CT 21-23 by the average hourly activity for CT 18-23. (C) Evening Anticipation Index (EAI) calculated by dividing the average hourly activity for CT 9-11 by the average hourly activity for CT 6-11. For scatter plots, each point represents an individual fly and mean ± SEM is shown. Significance determined by one-way ANOVA followed by Tukey’s multiple comparisons test (for normally distributed samples; E and H) or Kruskal-Wallis nonparametric ANOVA (to account for non-normality of samples; F and G) followed by Dunn’s multiple comparisons test; reported p-values are multiplicity adjusted to account for multiple comparisons. *** = p < 0.001, n.s. = not significant p > 0.05.

**Supplementary Table 1.** Percent rhythmicity values using Lomb-Scargle periodogram automated analysis. Individual flies were classified as rhythmic if difference between amplitude and the value of the best fit line was greater >150, using a p<0.001 statistical cutoff.

| Driver | *tim-Gal4* |  | *repo-Gal4* |  | *Mai179-Gal4* |  | *Pdf-Gal4* |  |
| --- | --- | --- | --- | --- | --- | --- | --- | --- |
| Scoring Method | Manual | Lomb-Scargle | Manual | Lomb-Scargle | Manual | Lomb-Scargle | Manual | Lomb-Scargle |
| *acp*<sup>CRISPR</sup> | 96.88 | 84.85 | 100.0 | 96.77 | 90.88 | 93.94 | 100.0 | 95.12 |
| *per*<sup>CRISPR</sup> | 0.00 | 0.00 | 96.77 | 93.55 | 0.00 | 3.13 | 74.19 | 86.67 |
| *tim*<sup>CRISPR</sup> | 0.00 | 2.70 | 96.88 | 87.50 | 0.00 | 3.28 | 83.33 | 91.67 |

| Driver | *Mai179-Gal4;*<br>*Pdf-Gal80* |  |
| --- | --- | --- |
| Scoring Method | Manual | Lomb-Scargle |
| *acp*<sup>CRISPR</sup> | 90.91 | 93 |
| *per*<sup>CRISPR</sup> | 86.04 | 94 |
| *tim*<sup>CRISPR</sup> | 84.62 | 89 |

**Supplementary Table 2.**
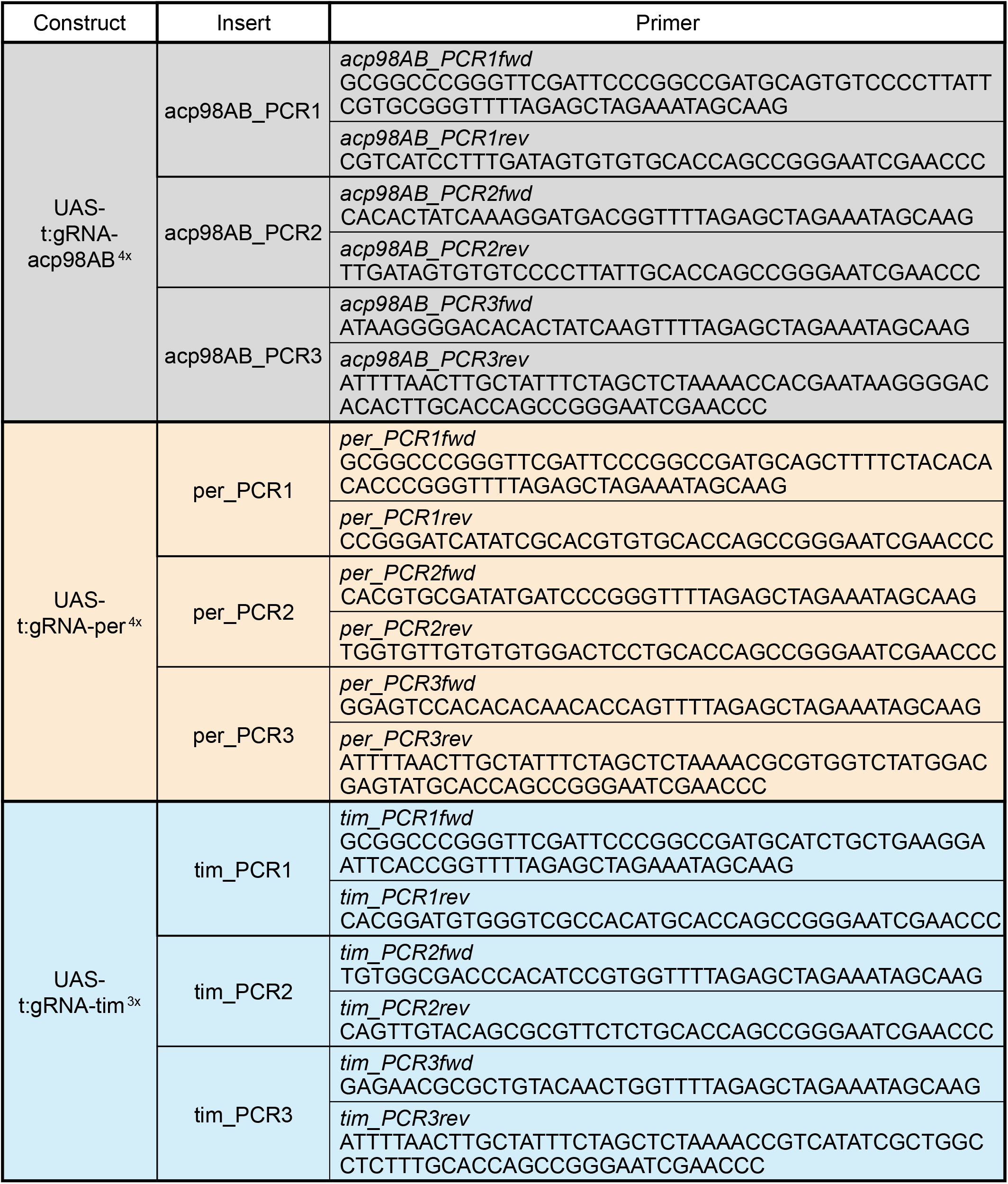
Primers to clone inserts for *UAS-sgRNA* construct assembly.

## References

1. Allen VW, et al. (2016) *period*-regulated feeding behavior and TOR signaling modulate survival of infection. Curr Biol 26(2):184–194.

2. Ulgherait M, et al. (2016) Dietary restriction extends the lifespan of circadian mutants *tim* and *per*. Cell Metab 24(6):763–764.

3. Stone EF, et al. (2012) The circadian clock protein timeless regulates phagocytosis of bacteria in *Drosophila*. PLoS Pathog 8(1):e1002445.

4. Hill VM, O’Connor RM, & Shirasu-Hiza M (2018) Tired and Stressed: examining the need for sleep. Eur J Neurosci.

5. Shirasu-Hiza MM, Dionne MS, Pham LN, Ayres JS, & Schneider DS (2007) Interactions between circadian rhythm and immunity in *Drosophila melanogaster*. Curr Biol 17(10):R353–355.

6. Panda S (2016) Circadian physiology of metabolism. Science 354(6315):1008–1015.

7. Vaccaro A, Issa AR, Seugnet L, Birman S, & Klarsfeld A (2017) *Drosophila* Clock Is Required in Brain Pacemaker Neurons to Prevent Premature Locomotor Aging Independently of Its Circadian Function. PLoS Genet 13(1):e1006507.

8. Rosbash M & Takahashi JS (2002) Circadian rhythms: the cancer connection. Nature 420(6914):373–374.

9. Maury E, Ramsey KM, & Bass J (2010) Circadian rhythms and metabolic syndrome: from experimental genetics to human disease. Circ Res 106(3):447–462.

10. Turek FW, Penev P, Zhang Y, van Reeth O, & Zee P (1995) Effects of age on the circadian system. Neurosci Biobehav Rev 19(1):53–58.

11. Wulff K, Gatti S, Wettstein JG, & Foster RG (2010) Sleep and circadian rhythm disruption in psychiatric and neurodegenerative disease. Nat Rev Neurosci 11(8):589–599.

12. Allada R, White NE, So WV, Hall JC, & Rosbash M (1998) A mutant *Drosophila* homolog of mammalian Clock disrupts circadian rhythms and transcription of *period* and *timeless*. Cell 93(5):791–804.

13. Hunter-Ensor M, Ousley A, & Sehgal A (1996) Regulation of the *Drosophila* protein timeless suggests a mechanism for resetting the circadian clock by light. Cell 84(5):677–685.

14. Rutila JE, et al. (1998) CYCLE is a second bHLH-PAS clock protein essential for circadian rhythmicity and transcription of *Drosophila period* and *timeless*. Cell 93(5):805–814.

15. Sehgal A, Price JL, Man B, & Young MW (1994) Loss of circadian behavioral rhythms and per RNA oscillations in the *Drosophila* mutant *timeless*. Science 263(5153):1603–1606.

16. Sehgal A, et al. (1995) Rhythmic expression of *timeless*: a basis for promoting circadian cycles in period gene autoregulation. Science 270(5237):808–810.

17. Vosshall LB, Price JL, Sehgal A, Saez L, & Young MW (1994) Block in nuclear localization of period protein by a second clock mutation, *timeless*. Science 263(5153):1606–1609.

18. Ko CH & Takahashi JS (2006) Molecular components of the mammalian circadian clock. Hum Mol Genet 15 Spec No 2:R271–277.

19. Top D & Young MW (2018) Coordination between differentially regulated circadian clocks generates rhythmic behavior. Cold Spring Harb Perspect Biol 10(7):1–27.

20. Stoleru D, Peng Y, Agosto J, & Rosbash M (2004) Coupled oscillators control morning and evening locomotor behaviour of *Drosophila*. Nature 431(7010):862–868.

21. Grima B, Chelot E, Xia R, & Rouyer F (2004) Morning and evening peaks of activity rely on different clock neurons of the *Drosophila* brain. Nature 431:869–873.

22. Yao Z, Bennett AJ, Clem JL, & Shafer OT (2016) The *Drosophila* clock neuron network features diverse coupling modes and requires network-wide coherence for robust circadian rhythms. Cell Rep 17(11):2873–2881.

23. Renn S, Park JH, Rosbash M, Hall JC, & Taghert PH (1999) A pdf neuropeptide gene mutation and ablation of PDF neurons each cause severe abnormalities of behavioral circadian rhythms in *Drosophila*. Cell 99:791–802.

24. Helfrich-Forster C (1995) The *period* clock gene is expressed in central nervous system neurons which also produce a neuropeptide that reveals the projections of circadian pacemaker cells within the brain of *Drosophila melanogaster*. Proc Natl Acad Sci U S A 92(2):612–616.

25. Fernandez MP, Berni J, & Ceriani MF (2008) Circadian remodeling of neuronal circuits involved in rhythmic behavior. PLoS Biol 6(3):e69.

26. Shafer OT, et al. (2008) Widespread receptivity to neuropeptide PDF throughout the neuronal circadian clock network of *Drosophila* revealed by real-time cyclic AMP imaging. Neuron 58(2):223–237.

27. Yao Z & Shafer OT (2014) The *Drosophila* circadian clock is a variably coupled network of multiple peptidergic units. Science 343(6178):1516–1520.

28. Martinek S & Young MW (2000) Specific genetic interference with behavioral rhythms in *Drosophila* by expression of inverted repeats. Genetics 156:1717–1725.

29. Shafer OT & Taghert PH (2009) RNA-interference knockdown of *Drosophila* pigment dispersing factor in neuronal subsets: the anatomical basis of a neuropeptide’s circadian functions. PLoS One 4(12):e8298.

30. Ng FS, Tangredi MM, & Jackson FR (2011) Glial cells physiologically modulate clock neurons and circadian behavior in a calcium-dependent manner. Curr Biol 21(8):625–634.

31. Yang Z & Sehgal A (2001) Role of molecular oscillations in generating behavioral rhythms in *Drosophila*. Neuron 29:453–467.

32. Gratz SJ, et al. (2013) Genome engineering of *Drosophila* with the CRISPR RNA-guided Cas9 nuclease. Genetics 194(4):1029–1035.

33. Yu Z, et al. (2013) Highly efficient genome modifications mediated by CRISPR/Cas9 in *Drosophila*. Genetics 195(1):289–291.

34. Port F, Chen HM, Lee T, & Bullock SL (2014) Optimized CRISPR/Cas tools for efficient germline and somatic genome engineering in *Drosophila*. Proc Natl Acad Sci U S A 111(29):E2967–2976.

35. Port F & Bullock SL (2016) Augmenting CRISPR applications in *Drosophila* with tRNA-flanked sgRNAs. Nat Methods 13(10):852–854.

36. Wolfner MF, et al. (1997) New genes for male accessory gland proteins in *Drosophila melanogaster*. Insect Biochem Mol Biol 27(10):825–834.

37. Gelbart WM, Emmert, D.B. (2013) FlyBase High Throughput Expression Pattern Data (FlyBase).

38. Kaneko M, Park JH, Cheng Y, Hardin PE, & Hall JC (2000) Disruption of synaptic transmission or clock-gene-product oscillations in circadian pacemaker cells of *Drosophila* cause abnormal behavioral rhythms. J Neurobiol 43(3):207–233.

39. Siegmund T & Korge G (2001) Innervation of the ring gland of *Drosophila melanogaster*. J Comp Neurol 431(4):481–491.

40. Park JH, et al. (2000) Differential regulation of circadian pacemaker output by separate clock genes in *Drosophila*. Proc Natl Acad Sci U S A 97(7):3608–3613.

41. Guo F, Cerullo I, Chen X, & Rosbash M (2014) PDF neuron firing phase-shifts key circadian activity neurons in *Drosophila*. Elife 3.

42. Konopka RJ & Benzer S (1971) Clock mutants of *Drosophila melanogaster*. Proc Natl Acad Sci U S A 68(9):2112–2116.

43. Young MW (1998) The molecular control of circadian behavioral rhythms and their entrainment in *Drosophila*. Annual Review of Biochemistry 67:135–152.

44. Hardin PE, Hall JC, & Rosbash M (1990) Feedback of the *Drosophila period* gene product on circadian cycling of its messenger RNA levels. Nature 343(6258):536–540.

45. Darlington TK, et al. (1998) Closing the circadian loop: CLOCK-induced transcription of its own inhibitors *per* and *tim*. Science 280(5369):1599–1603.

46. Price JL, Dembinska ME, Young MW, & Rosbash M (1995) Suppression of PERIOD protein abundance and circadian cycling by the *Drosophila* clock mutation timeless. EMBO J 14(16):4044–4049.

47. Price JL, et al. (1998) double-time is a novel *Drosophila* clock gene that regulates PERIOD protein accumulation. Cell 94(1):83–95.

48. Halter DA, et al. (1995) The homeobox gene *repo* is required for the differentiation and maintenance of glia function in the embryonic nervous system of *Drosophila melanogaster*. Development 121(2):317–332.

49. Xiong WC, Okano H, Patel NH, Blendy JA, & Montell C (1994) *repo* encodes a glial-specific homeo domain protein required in the *Drosophila* nervous system. Genes Dev 8(8):981–994.

50. Rieger D, Wulbeck C, Rouyer F, & Helfrich-Forster C (2009) *Period* gene expression in four neurons is sufficient for rhythmic activity of *Drosophila melanogaster* under dim light conditions. J Biol Rhythms 24(4):271–282.

51. Lin Y, Stormo GD, & Taghert PH (2004) The neuropeptide pigment-dispersing factor coordinates pacemaker interactions in the *Drosophila* circadian system. J Neurosci 24(36):7951–7957.

52. Stoleru D, Peng Y, Nawathean P, & Rosbash M (2005) A resetting signal between *Drosophila* pacemakers synchronizes morning and evening activity. Nature 438(7065):238–242.

53. Zhang L, et al. (2010) DN1(p) circadian neurons coordinate acute light and PDF inputs to produce robust daily behavior in *Drosophila*. Curr Biol 20(7):591–599.

54. Bulthuis N, Spontak KR, Kleeman B, & Cavanaugh DJ (2019) Neuronal Activity in Non-LNv Clock Cells Is Required to Produce Free-Running Rest:Activity Rhythms in *Drosophila*. J Biol Rhythms:748730419841468.

55. Top D, Harms E, Syed S, Adams EL, & Saez L (2016) GSK-3 and CK2 Kinases Converge on Timeless to Regulate the Master Clock. Cell Rep 16(2):357–367.

56. Top D, et al. (2018) CK1/Doubletime activity delays transcription activation in the circadian clock. Elife 7.

57. Picot M, Cusumano P, Klarsfeld A, Ueda R, & Rouyer F (2007) Light activates output from evening neurons and inhibits output from morning neurons in the *Drosophila* circadian clock. PLoS Biol 5(11):e315.

58. Montague TG, Cruz JM, Gagnon JA, Church GM, & Valen E (2014) CHOPCHOP: a CRISPR/Cas9 and TALEN web tool for genome editing. Nucleic Acids Res 42(Web Server issue):W401–407.

59. Pfeiffer BD, et al. (2010) Refinement of tools for targeted gene expression in *Drosophila*. Genetics 186(2):735–755.

60. Liu X, et al. (1992) The *period* gene encodes a predominantly nuclear protein in adult *Drosophila*. J Neurosci 12(7):2735–2744.

61. Schindelin J, et al. (2012) Fiji: an open-source platform for biological-image analysis. Nat Methods 9(7):676–682.

